# Molecular rotors provide insight into the mechanism of formation and conversion of *α*-synuclein aggregates

**DOI:** 10.1101/2024.09.13.612428

**Authors:** Siân C. Allerton, Marina K. Kuimova, Francesco A. Aprile

## Abstract

*α*-Synuclein is an intrinsically disordered protein that forms amyloids in Parkinson’s disease. Currently, detection methods predominantly report on the formation of mature amyloids but have poor sensitivity to the early-stage, toxic oligomers. Molecular rotors are fluorophores that sense changes in the viscosity of their local environment. Here, we monitor *α* -synuclein oligomer formation using the fluorescence lifetime of molecular rotors. We detect oligomer formation and conversion into amyloids for *wild type* and two *α* -synuclein variants; the pathological mutant A30P and ΔP *α* -synuclein, which lacks a master regulator region of aggregation (residues 36-42). We report that A30P *α* -synuclein shows a similar rate of oligomer formation compared to *wild type α* -synuclein, whereas ΔP *α* -synuclein shows delayed oligomer formation. Additionally, both variants demonstrate a slower conversion of oligomers to amyloids. Our method provides a quantitative approach to unveiling the complex mechanism of *α* -synuclein aggregation which is key to understanding the pathology of Parkinson’s disease.

## INTRODUCTION

Second only to Alzheimer’s disease, Parkinson’s disease (PD) is the most prevalent neurodegenerative age-associated disorder.^1–3^ PD is characterized by the formation of intraneuronal inclusions, the Lewy bodies (LB), which are predominantly composed of aggregates of the intrinsically disordered protein *α*-synuclein (*α*Syn).^1,4–9^ The majority of these aggregates, called amyloids, have a fibrillar shape and are enriched *β*-sheets.^2,3, 10^ Although *α*Syn plays a role in the pathology of PD, this 4.5 kDa protein, located at the presynaptic terminals of neurons,^11,12^ plays a key role in multiple physiological processes including neurotransmitter synthesis and release, and synaptic vesicle recycling.^13–16^

Recent studies indicate that oligomers are more toxic than amyloid fibrils and may be responsible for the disease onset and progression.^3,5,17^ These oligomers are highly heterogeneous in terms of conformation and toxic mechanisms.

Currently, the standard method for monitoring the aggregation of *α*Syn is by fluorescence intensity assays using switch-on dyes such as Thioflavin T (ThT) and PROTEOSTAT.^8–20^ However, this method predominantly reports on the formation of mature amyloid fibrils and is poorly sensitive to oligomers, unless super resolution techniques are employed.^17, 18,21^ Fluorescence Lifetime Imaging Microscopy (FLIM) can monitor the distribution of decay time in a spatially and time-resolved manner, therefore enabling the detection of photophysical events which fluorescence intensity imaging cannot achieve.

We set out to investigate whether oligomer detection can be achieved using molecular rotors (MRs).^18,22^ MRs display high non-radiative decay rates upon excitation in low-viscosity environments due to unrestricted rotation about an intramolecular bond. In contrast, in more viscous or crowded environments, the rotation is restricted causing a decrease in the non-radiative decay rate. This results in high fluorescence intensity and long fluorescence lifetimes, the latter is particularly suited to concentration-independent monitoring.^18,22,23^ In a crowded protein solution, the fluorescence lifetime of a MR can change to reflect the microviscosity of the environment, or due to crowding of MR in the presence of different structural species formed during the aggregation.

We previously demonstrated that the MRs, ThT and DiSC_2_ (3,3’-diethylthiacarbocyanine iodide) were able to follow the aggregation pathway of lysozyme and insulin, since aggregation caused the solution free volume sensed by these rotors to reduce, resulting in longer fluorescence lifetimes. Interestingly, the aggregation trajectory sensed by the lifetimes of ThT and DiSC_2_ were complex, indicating that the rotor-based detection was sensitive to species other than just the monomers and fully formed fibrils.^18^

Here, we explore the exciting potential of MR intensity and lifetime-based techniques to reveal the complexities of the aggregation mechanisms for *α*Syn involving oligomeric species. This is particularly important as establishing methods to identify and monitor the formation of oligomers is key to developing therapeutic approaches to target these potentially toxic aggregates, aiding in the development of clinical strategies for PD.

## RESULTS

### The fluorescence lifetime of DiSC_2_ and ThT increases before intensity during the aggregation of WT *α*Syn

The time-resolved fluorescence decays of DiSC_2_ and ThT were measured during the aggregation of *wild-type* (WT) *α*Syn, alongside traditional intensity measurements (Fig. 1). We selected ThT and DiSC_2_ as our MRs as both dyes are known to interact with amyloid fibrils.^18,24,25^ The aggregations were carried out in a glass-bottomed 96-well plate which was agitated (600 rpm) in a bench top incubator at 37 °C. At ∼ 30 min intervals the intensity (using a plate reader) and the time-resolved fluorescence decays (using a FLIM microscope) were recorded.

**Fig 1.**
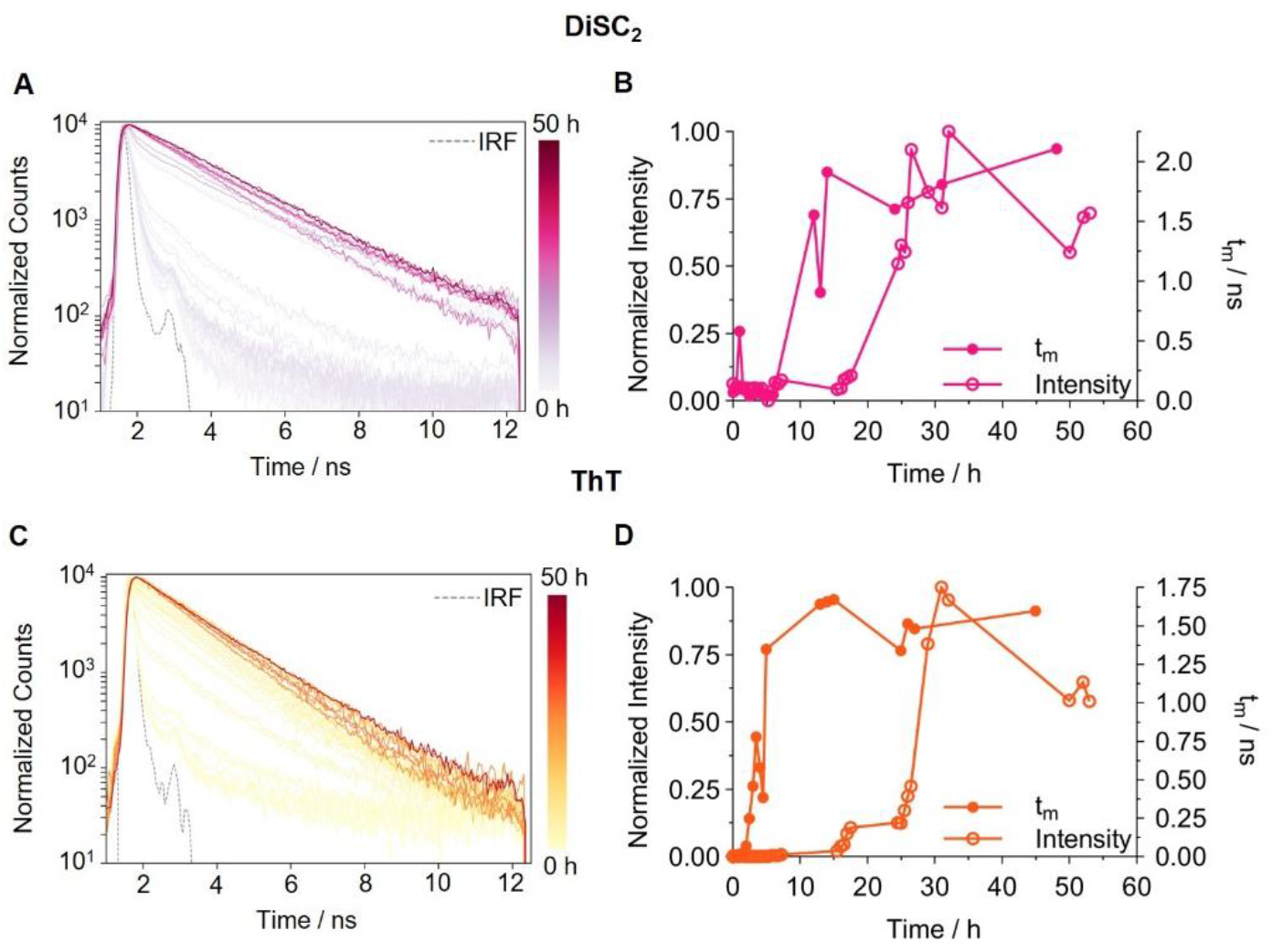
Time-resolved fluorescence decays and *τ*_m_ analysis of DiSC_2_ and ThT monitored during WT *α*Syn aggregation. **A** Single representative set of time-resolved fluorescence decays of DiSC_2_ (3 *μ*M) recorded during WT *α*Syn (50 *μ*M) aggregation. **B** Comparative studies of the intensity (hollow circles) and *τ*_m_ (filled circles) calculated from the DiSC_2_ time-resolved decays in Fig. A. **C** Single representative set of time-resolved fluorescence decays of ThT (0 *μ*M) recorded during WT *α*Syn (50 *μ*M) aggregation. **D** Comparative studies of the intensity (hollow circles) and *τ* _m_ (filled circles) calculated from the ThT time-resolved decays in Fig. C. The full datasets from all repeats are shown in Supplementary Fig. S4 (DiSC_2_) and S5 (ThT).

The fluorescence decays of DiSC_2_ and ThT (Fig. 1A, C) are initially dominated by a short-time component. This is indicative of the MR being in a non-viscous environment, most likely free in solution with purely monomeric WT *α*Syn, which does not restrict the MRs rotation. As the aggregation proceeds, the fluorescence lifetime of both rotors increases. This was expected as the aggregating species cause higher crowding which results in increased confinement of the MRs. Alternatively, the MR may bind to certain type(s) of aggregates, restricting its non-radiative relaxation to ground state in the bound conformation. However, if this was the case, we would expect to detect biexponential decays with fixed components and varying amplitudes, due to unbound (short component) and bound (long component) conformers of the MR. This is not the case and therefore we can exclude a simple binding model where MRs bind to a single species present in the aggregation mixture, *i*.*e*. WT *α*Syn mature fibrils.

Both MRs exhibit a similar rise in fluorescence intensity, after ∼ 15 h, indicating the intensity of both is sensitive to the presence of mature fibrils, consistent with literature data for ThT.^19,26^ Interestingly, the amplitude-weighted average lifetime (*τ*_m_) of DiSC_2_ and ThT, plotted against aggregation time reveals a much earlier increase compared to fluorescence intensity (Fig. 1B, D).

An increase in ThT *τ*_m_ is observed after 2 h of aggregation whereas the DiSC_2_ *τ*_m_ increases only after 12 h (Fig.1B, D). This data suggests that while the lifetime for both MRs can detect earlier formed species relative to intensity, ThT is more sensitive to these species compared to DiSC_2_. Alternatively, the two MRs may be sensitive to structurally distinct species.

The decay profiles of DiSC_2_ and ThT are complex, requiring overall 2 or 3 decay components to fit the time-resolved traces adequately. While the *τ*_m_ values reveal an overall increase in viscosity of the aggregating sample and can help identify kinetic change, the species causing these changes cannot be determined. Neither can we rule out that very low concentrations of fibrillar species are causing these changes. To identify these species’ and distinguish an increase in *τ*_m_ due to oligomers or fibrils, phasor analysis was used.

### The fluorescence lifetime of DiSC_2_ and ThT is sensitive to early-stage aggregates

Phasor plots represent a Fourier transform-based analysis which obtains a real (g) and imaginary (s) component of each fluorescence time-resolved decay associated with each time point during an aggregation. This is a non-biased analysis that does not require assumptions on the decay model, but instead the shape of the phasor trajectory during aggregation can help assign the nature of different species detected. Points in a similar position on the universal semicircle likely correspond to a similar environment (monomeric, oligomeric and fibrillar) and points lying along a straight line are likely from the same environments or species but at a varying concentration.^18,27^

The individual phasor plots of DiSC_2_ and ThT from multiple repeats of monitoring WT *α*Syn aggregation were combined to create a full phasor plot (Fig. 2A, C). In general, the phasor plots show a progression of points from the bottom right to the upper left of the universal semicircle, consistent with increased fluorescence lifetimes. Alongside aggregation, fibril phasor plots are shown in black, where increasing concentrations (0.00 -50 *μ*M) of pre-formed fibrils were measured in the presence of a fixed concentration of MRs (3 *μ*M DiSC_2_ and 0 *μ*M ThT). These phasor points represent confinement conditions created by fibrils alone. Thus, any differences between aggregation and fibril points indicate the detection of non-fibrillar species. However, the similarities in phasor points might mean either (i) that the points belong to fibrils present in the aggregation mixture, or (ii) that they coincide due to a similar environment created by oligomers and fibrils.

**Fig 2.**
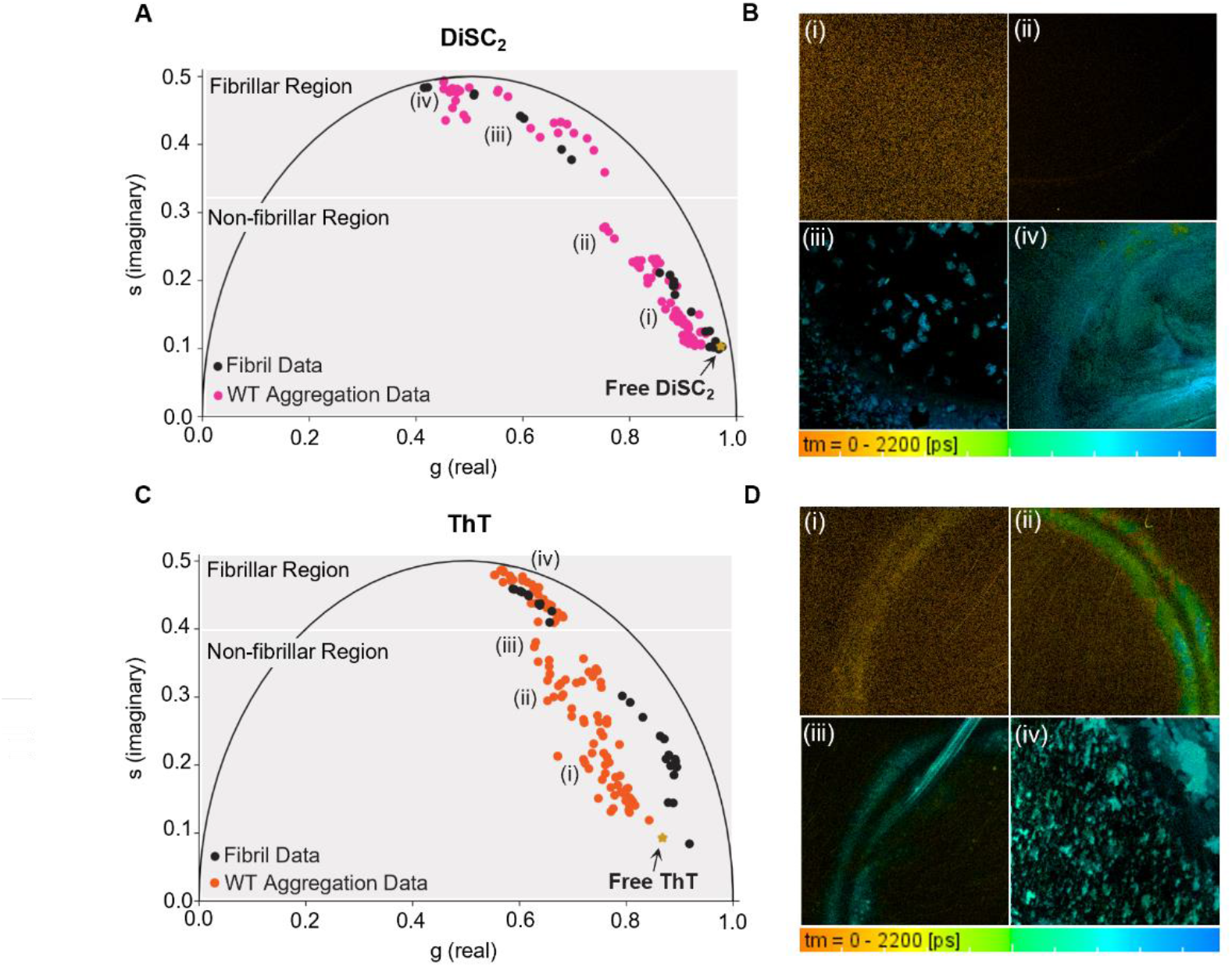
Phasor analysis and FLIM images of WT *α*Syn aggregation monitored with DiSC_2_ and ThT. **A** Phasor analysis of DiSC_2_ (3 *μ*M) in the presence of aggregating WT *α*Syn (50 *μ*M) (pink), data combined from four DiSC_2_ repeats (Supplementary Fig. S4). Phasor analysis of DiSC_2_ (3 *μ*M) in the presence of WT *α*Syn fibrils (0.00 -50 *μ*M, black) is overlayed (Supplementary Fig. S6). Any differences between the aggregation and the fibril plots indicates the presence and detection of non-fibrillar species in the aggregating solution. The fibrillar and oligomeric region highlighted in gray represent the regions in which the MRs are detecting specific species during the aggregation. Selection of the fibrillar and non-fibrillar regions (shown in gray) was based on the degree of overlap between the aggregation data and the fibrillar plot. Substantial overlap is considered when at least 60 % of aggregation points in that region overlap with the fibrillar data. Importantly, they do not represent the only regions where these species reside on the phasor plot (See Supplementary Fig. S 9 for further description). **B** FLIM images of DiSC_2_ associated with the aggregation data set shown in Fig. A, B. The images were recorded at (i) 3 h, (ii) 5.5 h, (iii) 2 h and (iv) 24 h. **C** Phasor analysis of ThT (0 *μ*M) in the presence of aggregating WT *α*Syn (50 *μ*M) (orange), data combined from five ThT repeats (Supplementary Fig. S5). Phasor analysis of ThT (0 *μ* M) in the presence of WT *α*Syn fibrils (0.00 -50 *μ*M, black) is additionally overlayed (Supplementary Fig. S7). Species specific regions, highlighted in gray, are assigned as described for DiSC_2_ (See Supplementary Fig. S20 for further description). **D** FLIM images of ThT associated with the aggregation data set shown in Fig. C, D. The images were recorded at (i) 2 h, (ii) 2.5 h, (iii) 3 h and (iv) 27 h. It should be noted that, for both DiSC_2_ and ThT, depending on the aggregation repeat slight differences in the time frame can be observed.

The DiSC_2_ phasor aggregation phasor plot (Fig. 2A) can be separated into two distinct regions. The first region (non-fibrillar region, Fig. 2A, B) occurs early in the aggregation when monomeric and smaller aggregates are expected to dominate and where we observe a partial deviation from the fibril trajectory. This suggests we are detecting the presence of a species with a structure that differs from mature fibrils. The second region (fibrillar region, Fig. 2A) occurs when aggregates at a size above the resolution limit of optical microscopy (300 nm) are observed in the lifetime images (Fig. 2B), suggesting that fibrillar species are present and dominate the signal of the MR. This is supported by phasor analysis of DiSC_2_ lifetimes in the presence of increasing fibril concentrations, which shows a clear overlap of phasor coordinates at later aggregation times.

The individual aggregation monitored by DiSC_2_ in Fig. 1B enters the fibrillar region of the phasor plot after 5.5 h (Fig. 2A (iii)), this is associated with a substantial *τ*_m_ increase (Fig. 1B). Interestingly, before 5.5 h, a small increase from 0.07 ns to 0.15 ns is observed (Fig. 1B), which is associated with a movement of phasor points in non-fibrillar region of the phasor plot (Fig. 2A). The *τ*_m_ of DiSC_2_, therefore, does increase in the presence of oligomeric species; however, the most substantial *τ*_m_ increase occurs when the formation of fibrils begins.

The aggregation of WT *α*Syn monitored by ThT (Fig. 2C, D) shows significant deviation from the fibril phasor plot in the early stages of the aggregation (non-fibrillar region, Fig. 2C) and overlap only occurs later in the pathway (fibrillar region, Fig. 2C). This suggests ThT is sensitive to an early aggregate species which has a distinctly different structure to mature fibrils, likely oligomers.^28^

The individual aggregation monitored by ThT in Fig. 1D enters the fibrillar region after 3 h (Fig. 2C (iv)), therefore the initial increase in ThT *τ*_m_ observed can be unequivocally attributed to the presence of oligomeric species.

### DiSC_2_ and ThT are sensitive to the presence of structurally distinct oligomers

It has been reported that the aggregation pathway of WT *α*Syn involves the formation of two types of oligomers, called Type-A and Type-B.^28,29^ Type-A oligomers can be considered an early-stage species whilst Type-B oligomers are a late-stage species which possess a higher *β*-sheet content compared to Type-A oligomers.^28^ We sought to determine whether the MRs were specific to either oligomer type.

To do this, stabilized oligomers, which have been shown to structurally resemble Type-B oligomers^17^, were prepared. In a similar manner to the fibril phasor plots, the lifetime decays of increasing concentrations of stabilized oligomers in the presence of a constant concentration of the MRs was measured.

The stabilized oligomers detected by DiSC_2_ reside at the bottom right of the phasor plot (Fig. 3A). These points show a partial overlap with the fibril phasor points, which can be due to the fact that both stabilized oligomers and fibrils have significant *β*-sheet content. Importantly, the early timepoints of WT *α*Syn aggregation show a much better overlap with the stabilized oligomer phasor points than with the fibril phasor points (Fig. 3B). We conclude that the early-stage aggregates sensed by DiSC_2_ are structurally similar to the stabilized oligomers, *i*.*e*., Type-B oligomers. We define the region of the phasor plot where the points from the aggregation of WT *α*Syn and the stabilized oligomers overlap as the *Type-B oligomeric region* (Fig. 3B). These results also suggest that the initial increase in *τ*_m_ described in Fig. 1 is probably due to the formation of Type-B oligomers.

**Fig 3.**
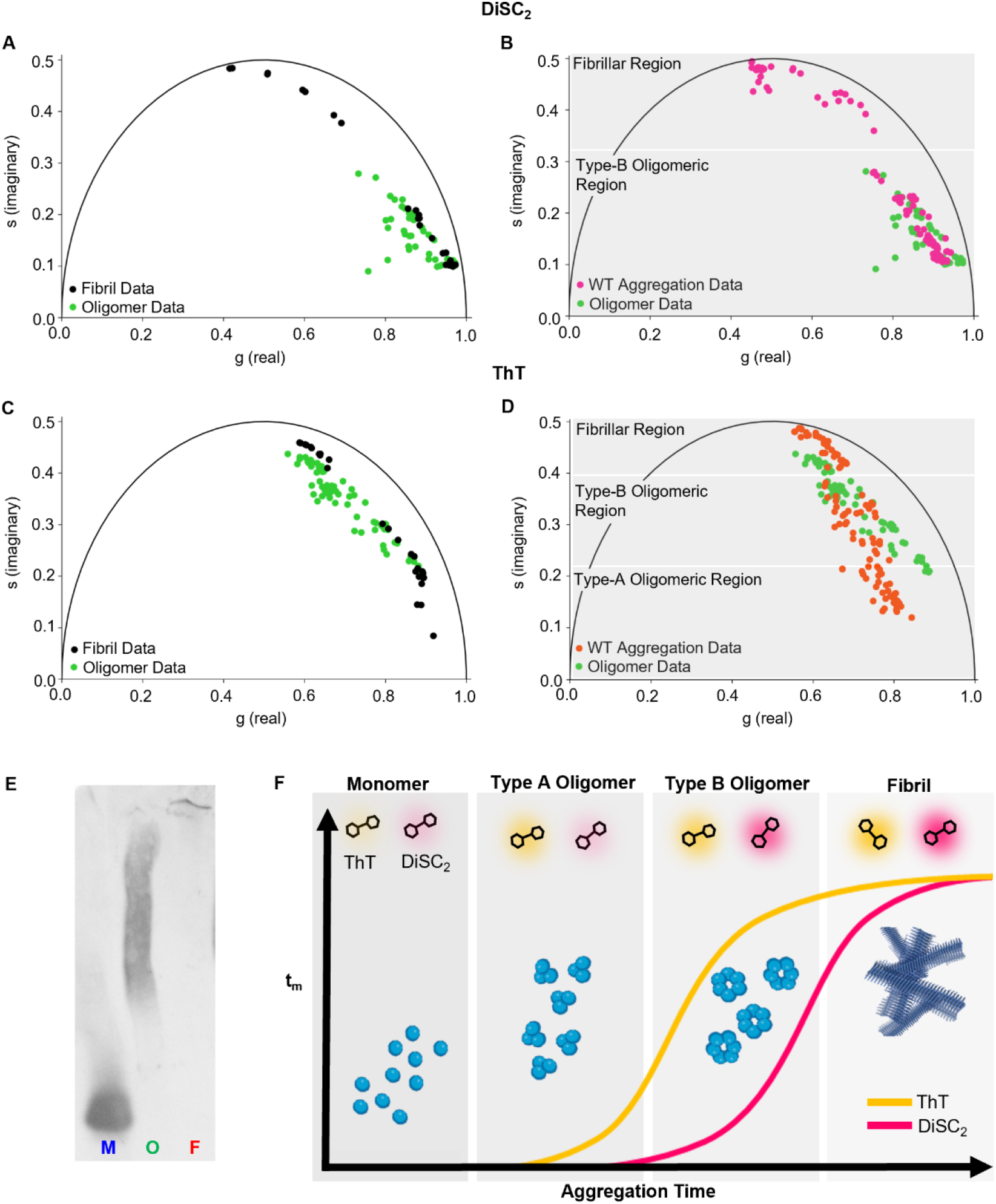
Comparative phasor analysis of time-resolved decay traces of DiSC_2_ and ThT the presence of different WT *α*Syn species. **A** Phasor analysis of DiSC_2_ (3 *μ*M) in the presence of isolated fibrils (black, 0.0 -50 *μ* M) and stabilized oligomers (green, 0.0 -20 *μ* M) (Supplementary Fig. S6, S8). **B** Phasor analysis of DiSC_2_ (3 *μ* M) in the presence of aggregating WT *α*Syn (50 *μ*M) (pink) as shown in Fig. 2A and stabilized oligomers (green, 0.0 -20 *μ*M) (Supplementary Fig. S4, S8). **C** Phasor analysis of ThT (0 *μ*M) in the presence of isolated fibrils (black, 0.0 -50 *μ* M) and stabilized oligomers (green, 0.0 -50 *μ* M) (Supplementary Fig. S7, S9). **D** Phasor analysis of ThT (0 *μ*M) in the presence of aggregating WT *α*Syn (50 *μ*M) (orange) as shown in Fig. 2C and stabilized oligomers (green, 0.0 -50 *μ*M) (Supplementary Fig. S5, S9). The fibrillar and oligomeric (Type-A and Type-B) regions highlighted in gray represent the regions in which the MRs are detecting specific species during the aggregation. Selection of the gray regions was based on the degree of overlap between the aggregation data, the fibrillar plot and stabilized oligomeric plot. Substantial overlap is considered when at least 60 % of aggregation points in that region overlap with the fibrillar or stabilized oligomer data. Importantly, they do not represent the only regions where these species reside on the phasor plot (See Supplementary Fig. S 9, S20 for further description). **E** Native-PAGE gel of monomeric (M), oligomeric (O) and fibrillar (F) WT *α*Syn. **F** Schematic diagram of the changes in *τ*_m_ of DiSC_2_ and ThT during the aggregation of WT *α*Syn, with the dominant species stated at each stage.

Similarly, we observed an overlap between WT *α* Syn aggregation phasor points and stabilized/Type-B oligomers detected by ThT (Fig. 3D). This suggests that also ThT senses aggregates which are structurally similar to stabilized oligomers, *i*.*e*., Type-B oligomers, and also in this case it is possible to define a phasor plot region as the *Type-B oligomeric region*. The fact that Type-B oligomers can be sensed by ThT explains the substantial *τ*_m_ increase, from 0.07 ns – 0.45 ns, observed between 2 and 3 h in Fig. 1D and Fig. 2C (i)-(iii). However, at very early timepoints, there is a clear deviation of the WT *α*Syn aggregation phasor plot from both the stabilized oligomer and fibril phasor points detected by ThT (Fig. 3D). These phasor points reflect an initial increase in *τ*_m_, from 0.012 to 0.7 ns, between 0 and 2 h in Fig. 1D and Fig. 2C (i). This observation suggests that ThT *τ*_m_ can initially sense the formation of aggregates which are structurally distinct from both stabilized oligomers and fibrils. This species may be Type-A oligomers, which are known to have a low *β*-sheet content,^17^ likely causing the large deviation from the other phasor plots. For this reason, we define this region of the phasor plot as the *Type-A oligomeric region*. Altogether, these results suggest that ThT can report on both the presence of Type-A and Type-B oligomers with the *τ*_m_ showing a large increase in the presence of Type-B oligomers compared to Type-A oligomers.

We have shown that DiSC_2_ and ThT can be used to monitor the aggregation of WT *α*Syn. Both MRs show an increase in *τ*_m_ which can be attributed to the presence of oligomers. Importantly, for both DiSC_2_ and ThT, phasor analysis can be utilized to identify the formation of oligomeric species (Type-A and Type-B for ThT and Type-B for DiSC_2_), even if the *τ*_m_ does not show a substantial increase.

### ThT lifetime confirms that the A30P *α*Syn variant has enhanced oligomerization

As shown above, ThT can report on both the presence of Type-A and -B oligomers while DiSC_2_ is sensitive to only Type-B (Fig. 2). ThT was therefore used to monitor the aggregation of two further variants which have been reported to have different aggregation kinetics with respect to WT *α*Syn. First, we investigated the behavior of the genetic mutant A30P *α*Syn, which has been reported to aggregate slower than WT *α* Syn and enable a build up of oligomers.

A substantial increase in ThT *τ*_m_ is observed after 2 h in the presence of aggregating WT and A30P *α*Syn (Fig. 4A) but the lag phase recorded by ThT intensity is extended for A30P *α*Syn compared to WT *α*Syn. This suggests that the rate of Type-B oligomer formation is similar for the two proteins, but the rate of oligomer conversion into fibrils is slower for A30P *α*Syn. Our results support previous studies which show that A30P *α*Syn has enhanced oligomerization compared to WT *α* Syn.^30,31^ The final lifetime values seen for A30P *α* Syn are also lower compared to WT *α*Syn. This can reflect a different structure of species present (both oligomers and fibrils), or a different mode of interaction of ThT with these protein species.

**Fig 4.**
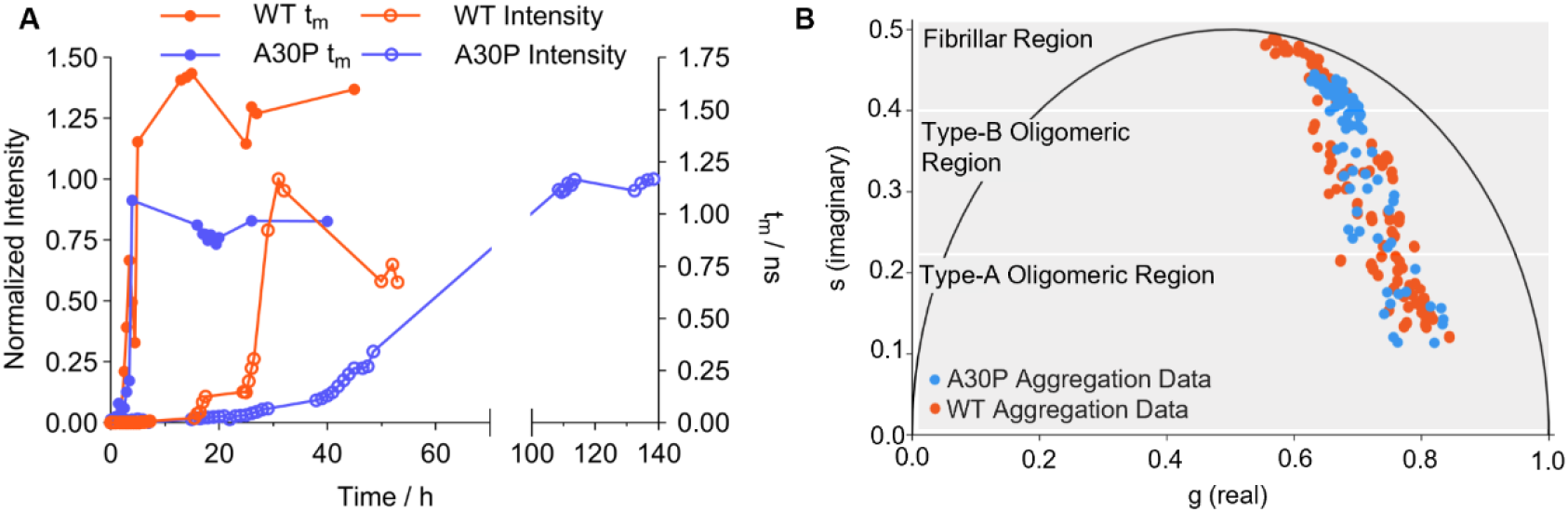
Lifetime analysis and comparative phasor plots of A30P and WT *α* Syn aggregation monitored with ThT. **A** Comparison of the intensity (hollow circle) and *τ*_m_ (full circle) of ThT (0 *μ*M) in the presence of A30P (blue, *μ*M) and WT *α*Syn (orange, *μ*M) during a single aggregation repeat, the full dataset from all repeats is shown in Supplementary Fig. S5, S 0. **B** Phasor analysis of the time-resolved decay traces of ThT (0 *μ*M) in the presence of aggregating A30P *α*Syn (blue, data combined from three repeats (Supplementary Fig. S 0)), overlayed with the data for WT *α*Syn (orange, data from Fig. 2).

There are small but significant differences between the aggregation phasors of WT and A30P *α*Syn involving time points collected both at early and late stages of aggregation (Fig. 4B). These differences indicate potential structural differences between the oligomers of A30P and WT *α*Syn. However, given that A30P *α*Syn has been reported to accumulate oligomers,^31^ we cannot exclude that these differences are due to a higher concentration of oligomers of A30P *α*Syn. In fact, increasing concentration of a protein species could cause a proportional shift in the lifetime.

### ΔP1 *α*Syn variant delays formation of oligomeric species during aggregation

The ‘P region’ of *α*Syn (residues 36-42) has recently been identified as a master regulator of amyloid aggregation.^32,33^ In fact, a variant of *α*Syn missing this protein region, *i*.*e*., ΔP1 *α*Syn, has been shown to accumulate as oligomers, resulting in reduced amyloid formation.^32,33^ We set out to test whether our method could uncover further details about the time evolution of the oligomers of this protein. We found that, for WT *α*Syn, ThT *τ*_m_ increased rapidly after 3 h, whereas for ΔP1 *α*Syn this increase was observed after ∼20 h (Fig. 5A). This observation suggests that the deletion of the P1 region delays the formation of oligomers. The increase in ThT intensity was also delayed, t_50_ ∼70 h for ΔP *α*Syn *vs* ∼30 h for WT *α*Syn (Fig. 5A). This suggests that the P region plays a key role in regulating the rate of both oligomer and fibril formation. Interestingly, deletion of the P region appears to cause a greater shift in the ThT intensity increase compared to the *τ*_m_ increase when compared to WT *α*Syn (+40 h *vs* + 7 h). This suggests that the conversion of oligomers to fibrils is more significantly impacted than the initial oligomer formation. This supports a recent study showing that the P region acts as a controller for the conversion of oligomers to fibrils.^34^

**Fig 5.**
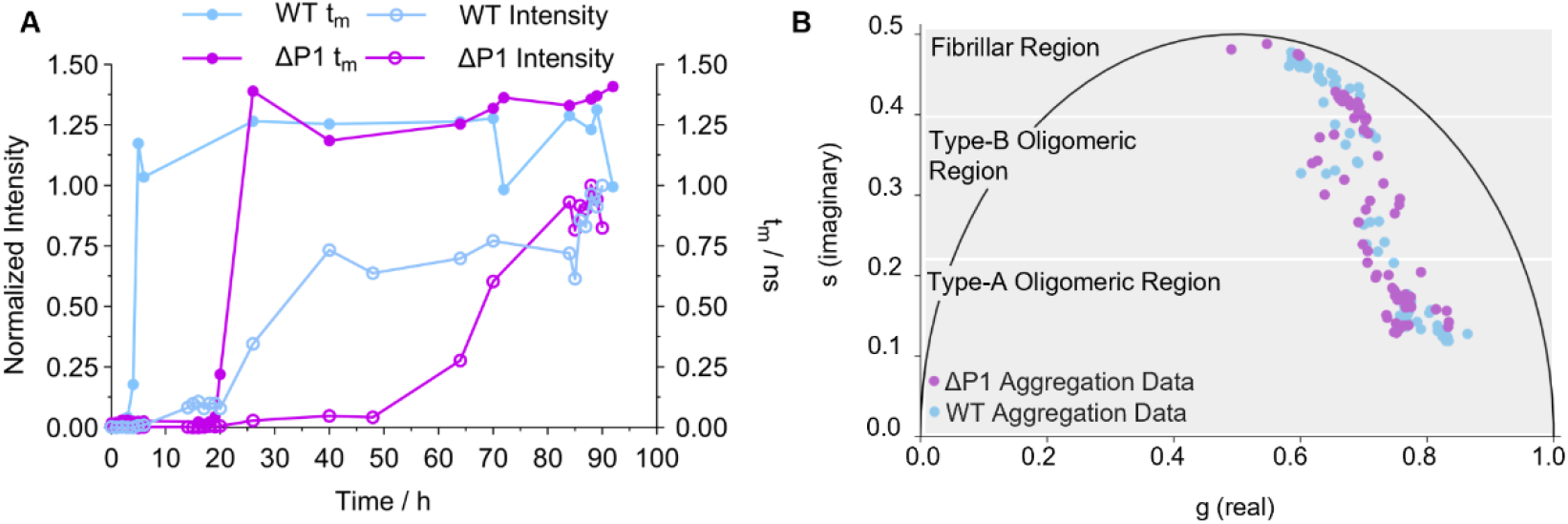
Lifetime analysis and comparative phasor plots of ΔP1 and WT *α*Syn aggregation monitored with ThT. **A** Comparison of the intensity (hollow circle) and *τ*_m_ (full circle) of ThT (0 *μ*M) in the presence of ΔP (purple, 00 *μ*M) and WT *α*Syn (light blue, 00 *μ*M) during a single aggregation repeat, the full dataset from all repeats is shown in Supplementary Figs. S 2, S 3. **B** Phasor analysis of the time-resolved decay traces of ThT (0 *μ*M) in the presence of aggregating ΔP *α*Syn (purple, data combined from three repeats (Supplementary Fig. S 3)), overlayed with the data from the aggregation of WT *α*Syn (light blue, data combined from three repeats (Supplementary Fig. 2).

The same final *τ*_m_ value (∼1.5 ns) (Fig. 5A) was reached for WT *α*Syn and ΔP1 *α*Syn, which indicates a similar degree of crowding for the dye in both variants and supports previous literature which suggests that the morphologies of the mature amyloid fibrils are similar.^32,34^ Furthermore, both WT and ΔP1 *α*Syn aggregation phasor trajectories show a good degree of overlap (Fig. 5B), suggesting similar structural interaction with ThT (*i*.*e*. similar proportions of species present) enroute to fibrils, albeit at very different kinetics (Fig. 5A).^32,34^

## CONCLUSIONS

We have developed a novel method based on the combined use of fluorescence lifetime and intensity of two molecular rotors, ThT and DiSC_2_,^18,24^ to monitor the aggregation of WT *α*Syn and its variants, in real time, including the formation of oligomers.

We found that the *τ*_m_ of both MRs was more sensitive to changes in the aggregation of WT *α*Syn compared to intensity (Fig. B, D). However, *τ*_m_ analysis alone could not help us identify the structural species present in the aggregating solution. For this, we utilized phasor analysis; a non-biased fitting analysis of individual time-resolved decays. We observed that the aggregation phasor plots of both MRs showed deviation from isolated fibril phasors indicating their ability to sense an early-stage aggregate, most likely oligomers (Fig. 2A, C).

To further characterize these early-stage aggregates we prepared stabilized oligomers which have been shown to be structurally like Type-B oligomers,^17^ a class of late-stage oligomers which have a *β*-sheet content similar to mature fibrils and are highly toxic.^28^ Type-A oligomers, the other main class of oligomers that form during an *in vitro α*Syn aggregation, form early in the aggregation and are considered weakly toxic.^28,29^

Phasor analysis of DiSC_2_ in the presence of stabilized oligomers showed a clear overlap with the early-stages of the aggregation phasor (Type-B oligomeric region, Fig. 3B). These data suggest that DiSC_2_ can be used to probe Type-B oligomers, however the *τ*_m_ only begins to significantly increase when the phasor enters the fibrillar region (Fig. 1B, 3B).

Phasor analysis of ThT in the presence of stabilized oligomers showed significant overlap with some of the aggregation phasor (Type-B oligomeric region, Fig. 3D), indicating ThT is also sensitive to the presence Type-B oligomers. Moreover, a key region of the phasor plot was identified where there is no overlap with the aggregation phasor points associated with the very early stages of aggregation and either the stabilized oligomer and fibril phasor plots (Type-A oligomeric region, Fig. 3D). Therefore, ThT can detect the presence of an early-stage oligomeric species which is structurally distinct from stabilized oligomers. These species are most likely Type-A oligomers. Importantly, a significant increase in *τ*_m_ is only observed when the ‘Type-B oligomeric region’ (Fig. 1D, 3D) is entered, indicating that the kinetic characterization of Type-B oligomers can be achieved using ThT.

Finally, we used our approach to monitor the oligomer formation of two variants of *α*Syn, named A30P and ΔP *α*Syn, using ThT. We validated our method on these two proteins because they present major differences in the aggregation behavior with respect to WT *α*Syn.^30,32,34^ For A30P *α*Syn, we found that ThT *τ*_m_ increases at a similar rate to that of WT *α*Syn during aggregation. This observation suggests that the rate of oligomer formation of A30P *α*Syn is comparable to that of the WT protein. Conversely, ThT intensity of A30P *α*Syn increases at a significantly lower rate. This result indicates that the oligomer conversion into fibrils of A30P *α*Syn is significantly delayed with respect to that of WT *α*Syn. The similar rate of formation of oligomer but delayed conversion of oligomers into fibrils could explain the accumulation of oligomers observed for A30P *α*Syn.^30,31^

In contrast, both ThT *τ*_m_ and intensity increases were delayed for ΔP *α*Syn compared to WT *α*Syn. This result suggests that the P region plays a role in regulating both oligomer and fibril formation. Nevertheless, the effect on ThT intensity was greater than that on ThT *τ*_m_, indicating that that deletion of the P region predominantly affects the conversion of oligomers into fibrils rather than the initial formation of the oligomers. These findings were in agreement with previous literature.^30,34^

Our novel molecular rotor-based approach provides an exciting new toolset for monitoring the aggregation of *α*Syn and provides structural insight into the aggregated species formed by the protein. The sensitivity of this approach for the oligomers demonstrates its high potential in comparison to standard bulk-averaged intensity measurements. By using fluorescence intensity and lifetime in combination, we can unveil the complicated mechanism of *α* Syn amyloid formation to identify and monitor key targets in PD pathology, therefore, opening doors to new diagnostic and therapeutic strategies.

## METHODS

### Expression and purification of WT *α*Syn

pT7-7 WT *α*Syn transformed BL2 -Gold (DE3) competent E. coli cells were used to express WT *α* Syn as per the manufacturer’s instructions (Agilent Technologies, Santa Clara, CA, USA). Expression was scaled up to 4 L and carried out at 37 °C for 4 h. To harvest the cells, the suspensions were centrifuged and resuspended in 20 mM Tris-HCl, mM EDTA, pH 8.0, including protease inhibitors. The subsequent purification of WT *α*Syn was then carried out as previously described.^35,36^ The lysate was sonicated on ice and the supernatant, following centrifugation (30 min, 8,000 rpm, 4 °C) was boiled at 80 °C for 20 min to precipitate heat-sensitive proteins. The denatured protein was removed by centrifugation (30 min, 8,000 rpm, 4 °C) and the supernatant was incubated with streptomycin sulfate, (20 mg/mL, 20 min, 4 °C) to precipitate the DNA which was subsequently removed by centrifugation (30 min, 8,000 rpm, 4 °C). The slow addition of ammonium sulfate (360 mg/mL, 20 min, 4 °C) while stirring precipitated out the *α* Syn which was collected by centrifugation and the pellet was resuspended in 20 mM Tris-HCl, mM EDTA, pH 8.0. To ensure complete buffer exchange, dialysis of the resuspended sample was carried out overnight at 4 °C. The protein solution was then purified by ion-exchange chromatography using a HiPrep Q HP 6/ 0 column (Cytiva Life Sciences) and the eluted protein was subsequently purified by size-exclusion chromatography using a HiLoad 6/600 Superdex 75 pg column (Cytiva Life Sciences). The purified protein was eluted in PBS (pH 7.4). The final purified protein concentration was determined by UV-Vis spectroscopy using a Cary 60 UV-Vis spectrometer (Agilent Technologies, Santa Clara, CA, USA) by recording the absorbance at 275 nm and a molar extinction coefficient of 5600 M^-1^ cm^-1^.

### Expression and purification of *α*Syn variants (A30P and ΔP1 *α*Syn)

The A30P *α*Syn plasmid was prepared by site-directed mutagenesis of the pT7-7 WT *α*Syn plasmid using a QuikChange II Site-Directed Mutagenesis Kit (Agilent, Agilent Technologies, Santa Clara, CA, USA), the primers, obtained from Eurofins Scientific, are given in the table below (Table 1).

**Table 1.**
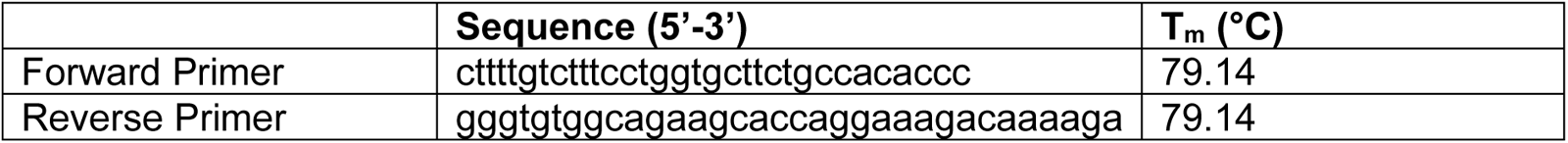
Sequence of the forward and reverse primer for the mutagenesis of pT7-7 WT *α*Syn plasmid to produce the A30P *α*Syn variant plasmid.

The ΔP *α*Syn plasmid was obtained from Professor Sheena E. Radford (University of Leeds). Both variants were then transformed and expressed following the same method as described above for WT *α* Syn. The concentration of the variants was determined by UV-Vis spectroscopy using a molar extinction coefficient of 5600 M^-1^ cm^-1^ for A30P *α*Syn and 4470 M^-1^ cm^-1^ for ΔP *α*Syn, at an absorbance at 275 nm. The variants were stored in PBS (pH 7.4). For the aggregations of ΔP *α*Syn the variant was buffer exchanged into 20 mM Tris-HCl with 200 mM NaCl (pH 7.5) or 20 mM sodium acetate with 200 mM NaCl (pH 4.5).

### Preparation of *α*Syn fibrils

*α*Syn fibrils were prepared by incubating high concentration solutions of monomeric *α*Syn (∼ 200 µM) at 37 °C for 5 days with agitation (∼ 600 rpm). The resulting fibrils were pelleted by centrifugation (4000 rpm) and resuspended in the required buffer. The centrifugation and resuspension steps were repeated three times. The final fibril concentration, following treatment with guanidinium hydrochloride (Gnd-HCl) to a final concentration of 4 M, was estimated by UV-Vis spectroscopy using a Cary 60 UV-Vis spectrometer (Agilent Technologies, Santa Clara, CA, USA) by recording the absorbance at 275 nm and a molar extinction coefficient of 5600 M^-1^ cm^-1^. This gave the concentration in monomer equivalents.

### Preparation of *α*Syn stabilized oligomers

*α*Syn stabilized oligomers were prepared using an adapted protocol from Chen et. al.^7^ 400 *μ* L of 800 *μ* M monomeric *α* Syn in PBS (pH 7.4) was flash frozen in liquid nitrogen and lyophilized overnight. The lyophilized protein was incubated at 37 °C for 24 h without agitation to promote the formation of oligomeric species. Fibrillar species were then removed by ultracentrifugation (55,000 rpm, .25 h) and the monomeric protein removed by multiple filtration steps using a 00 kDa cut-off membrane. The final oligomeric solution concentration was estimated by UV-Vis spectroscopy using a nanodrop at 275 nm and a molar extinction coefficient of 7000 M^-1^ cm^-1^.

### Amyloid formation monitored by fluorescence intensity and fluorescence lifetime imaging (FLIM)

WT and A30P *α*Syn (50 *μ*M) monomer solutions were aggregated in the presence of 0 *μ*M ThT or 3 *μ*M DiSC_2_, 0.02% sodium azide (NaN_3_) and PBS buffer (pH 7.4). 20 *μ*L of each sample was incubated in a 96 well full-area glass bottom plate (Sensoplate, Greiner Bio One, UK) with a borosilicate bead (3 mm diameter) at 37 °C for 24-48 h in a benchtop incubator, shaking at 450 rpm. The plate was covered with a clear film (ibiSeal, Ibidi, Germany). At ∼ 30 min intervals, the intensity and fluorescence lifetime was measured.

ΔP and WT *α*Syn (00 *μ*M) monomer solutions were aggregated in the presence of 0 *μ*M ThT, 0.02% sodium NaN_3_ and 20 mM Tris-HCl with 200 mM NaCl (pH 7.5). 00 *μ*L of each sample was incubated in a 96 well full-area glass bottomed plate (Sensoplate, Greiner Bio One, UK) at 37 °C for ∼ 60 h in a benchtop incubator, shaking at 600 rpm. The plate was covered with a clear film (ibiSeal, Ibidi, Germany). At ∼ 30 min intervals, the intensity and fluorescence lifetime were measured.

## Fluorescence intensity measurements

Fluorescence intensity measurements of the same samples measured by FLIM in the 96 well full-area glass bottomed plate (Sensoplate, Greiner Bio One, UK) was taken in a CLARIOstar Plus microplate reader (BMG Labtech, Ortenberg, Germany). Spiral averaging (3 mm diameter), excitation 440 nm, dichroic 460 nm, and emission 480 nm filters (ThT) or excitation 530 nm, dichroic 552.5 nm and emission 580 nm (DiSC_2_), 4 gains and 50 flashes per well was used. Aluminium foil was used on top of the clear film to cover the samples for the intensity measurements.

### Fluorescence lifetime imaging

The fluorescence lifetime images of samples on a borosilicate glass slide or 96 well full-area glass bottomed plate (Sensoplate, Greiner Bio One, UK) were obtained using a Leica TSC SP5 II inverted confocal microscope (Leica Microsystems GmbH, Germany) and a 0x air objective. A two-photon pulsed excitation (*λ*_ex_ = 880 nm for ThT and *λ*_ex_ = 950 nm for DiSC_2_) was provided by an external Ti:Sapphire laser (Chameleon Vision II, 80 MHz). Emission was recorded from 465 nm to 540 nm for ThT and from 525 nm to 700 nm for DiSC_2_. For each sample a fluorescence lifetime image at at least two z-positions was measured.

Fluorescence lifetime data analysis

The equation below (Equation) was used to fit the fluorescence decays to a mono-, bi-or tri-exponential model.

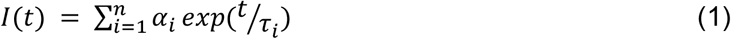

Where *I* is fluorescence intensity, *t* is time, and *τ*_*i*_ and *α*_*i*_ are the fluorescence lifetimes and amplitudes of the *n* exponentially decaying components, respectively. SPCImage (Becker & Hickl GmbH, Germany) was used to achieve the fitting. For the time-resolved decays that fit the bi-or triexponential models, the amplitude-weighted mean lifetime (*τ*_*m*_) was also determined following the equation below (Equation 2).

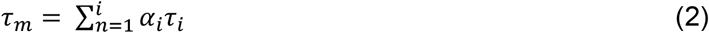

GraphPad Prism version 9.3. for Windows (GraphPad Software, San Diego, CA, USA) was used to plot the data. Phasor analysis was achieved using software written in-house in Matlab (MathWorks) and was performed on the fluorescence lifetime decays of the monomers and aggregates of WT, A30P and ΔP *α*Syn. By plotting the real (g) and imaginary (s) components of the Fourier transform of a fluorescence decay, a two-dimensional representation of the data can be obtained. The two components are calculated as shown below.

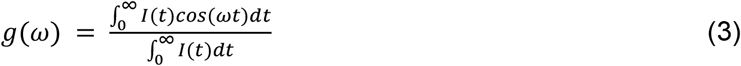

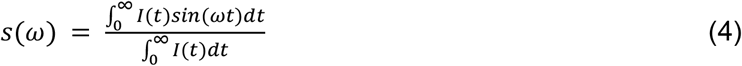

Where *ω* is the angular repetition frequency of the pulsed excitation laser (80 MHz), *I*(*t*) is the measured fluorescence decay and *t* is time. The phasors of the decays can be superimposed on a “universal circle” (central coordinates (/2, 0)). For homogenous systems in which only one fluorescence lifetime, *τ*, exists, the phasor points *g*(*ω*) = 1/[1 + (*ωτ*)^2^] (real) and *s*(*ω*) = *ωτ*/[1 + (*ωτ*)^2^] (imaginary) lie on the universal circle. This type of decay can be considered monoexponential, whereas phasors of multiexponential decays will lie within this circle.^8^ Phasor analysis can therefore give an indication of the number of different species which form during the aggregation of *α*Syn.

## ACKNOWLEDGEMENTS

The authors thank UK Research and Innovation (Future Leaders Fellowship MR/S033947/ and MR/Y0036 6/), the Engineering and Physical Sciences Research Council (Grant EP/S0235 8/), and Alzheimer’s Research UK (ARUK-PG20 9B-020) for support. We thank Professor Sheena E. Radford (University of Leeds) for providing us with the expression plasmid of the variant ΔP *α*Syn and for the helpful feedback on the manuscript.

