## Supplementary Information for "Molecular rotors provide insight into the mechanism of formation and conversion of *α*-synuclein aggregates"

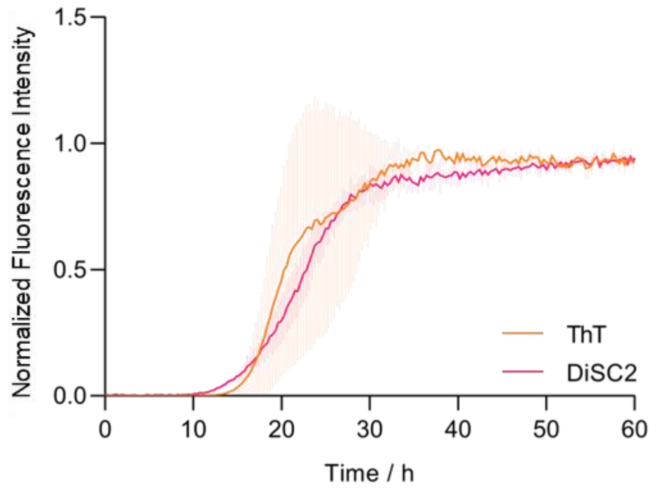


**Supplementary Fig. S1 Aggregation of WT $\alpha$Syn.** Fluorescence intensity assay of WT $\alpha$Syn (150 $\mu$M) in PBS (pH 7.4), monitored by $DiSC_2$ (3 $\mu$M) or ThT (10 $\mu$M). Done in triplicate.

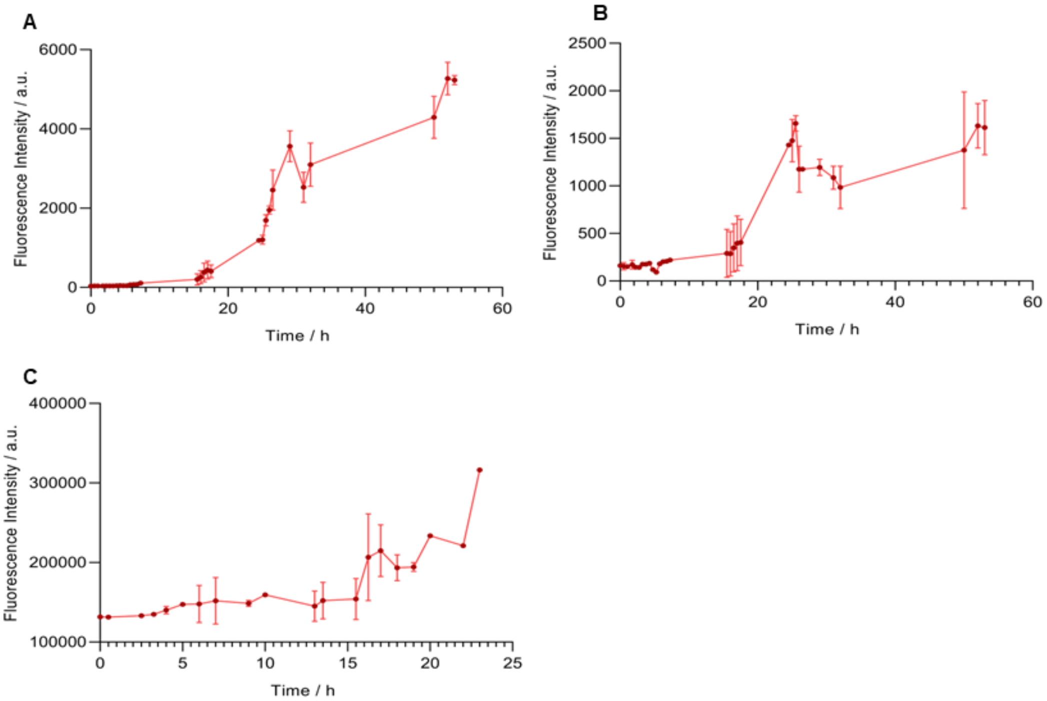


**Supplementary Fig. S2 Aggregation of WT and A30P $\alpha$Syn. A** Fluorescence intensity assay of WT $\alpha$Syn (150 $\mu$M) monitored by ThT (10 $\mu$M). **B** Fluorescence intensity assay of WT $\alpha$Syn (150 $\mu$M) monitored by $DiSC_2$ (3 $\mu$M). **C** Fluorescence intensity assay of A30P $\alpha$Syn (150 $\mu$M) monitored by ThT (10 $\mu$M). Done in duplicate.

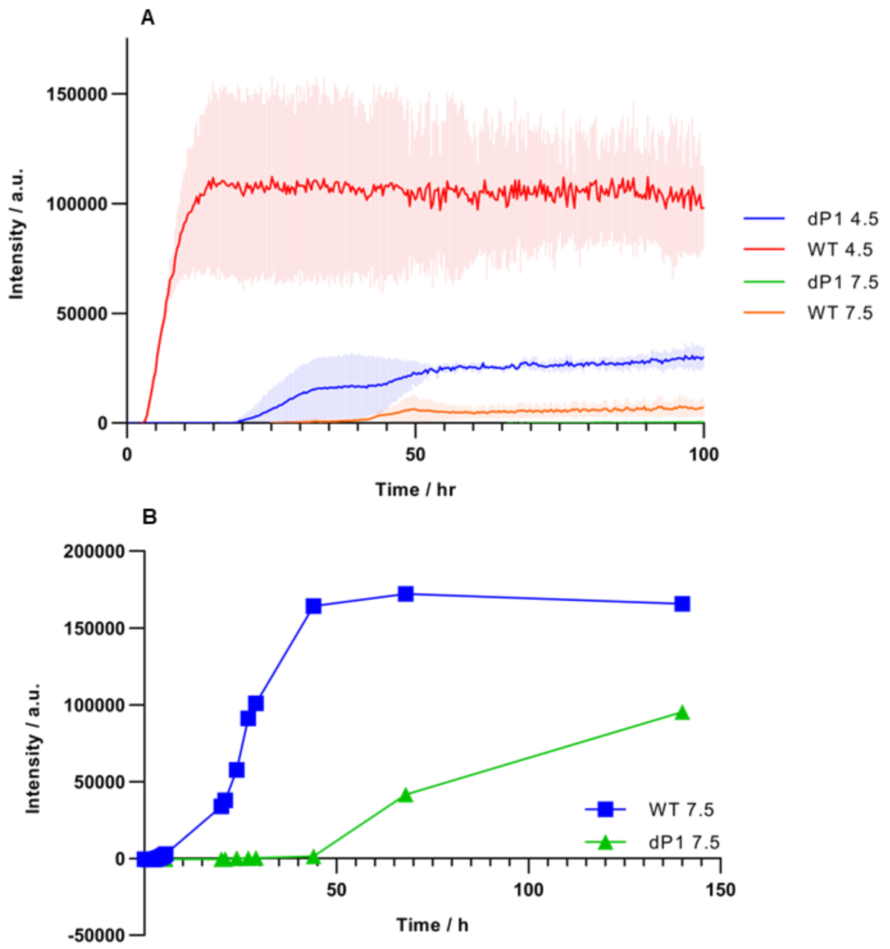


**Supplementary Fig. S3 Aggregation of WT and ΔP1 $\alpha$Syn. A** Fluorescence intensity assay of WT and ΔP1 αSyn (100 $\mu$M) in 20 mM Tris-HCl with 200 mM NaCl (pH 7.5) or 20 mM sodium acetate with 200 mM NaCl (pH 4.5), monitored by ThT (10 $\mu$M). Done in triplicate. **B** Fluorescence intensity assay of WT and ΔP1 αSyn (100 $\mu$M) in 20 mM Tris-HCl with 200 mM NaCl (pH 7.5) monitored by ThT (10 $\mu$M).

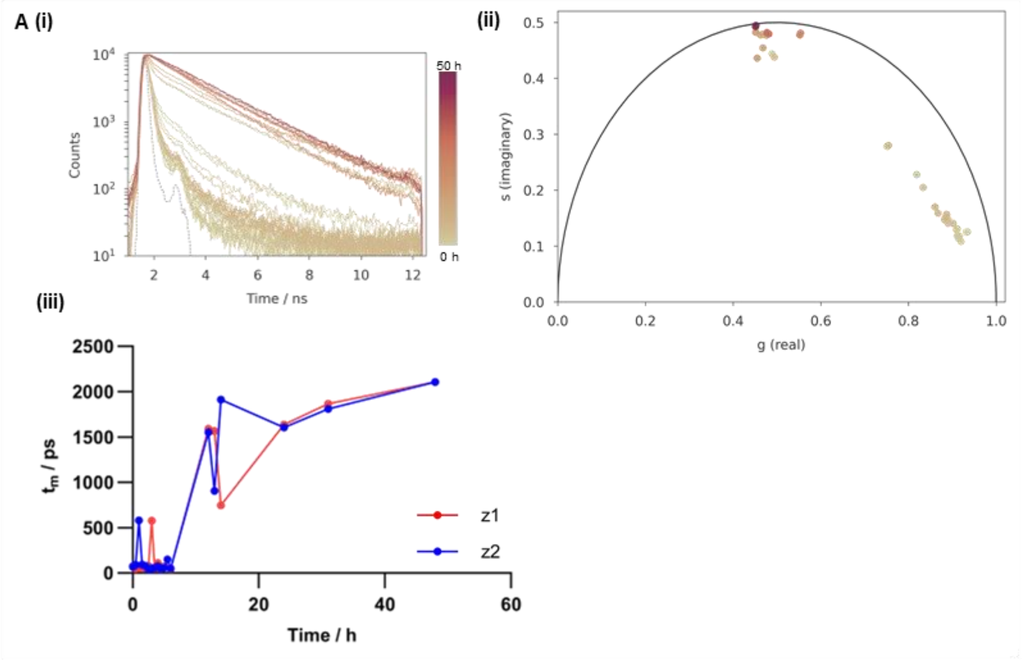
A (i)
(ii)
(iii)
Counts
Time / ns
50 h
0 h
s (imaginary)
g (real)
$t_m$ / ps
Time / h
z1
z2

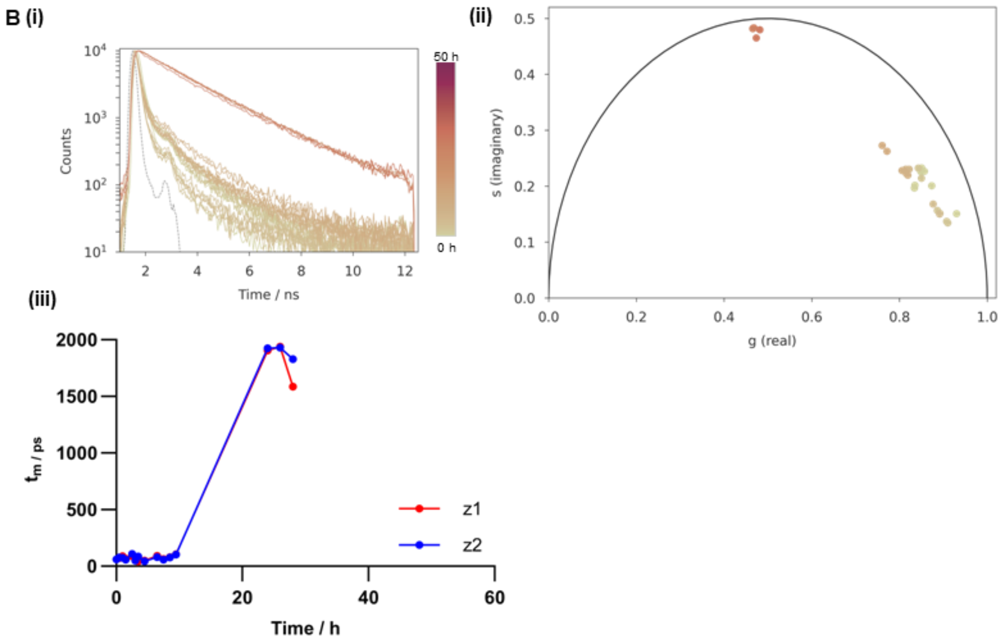
B (i)
(ii)
(iii)
Counts
Time / ns
50 h
0 h
s (imaginary)
g (real)
$t_m$ / ps
Time / h
z1
z2

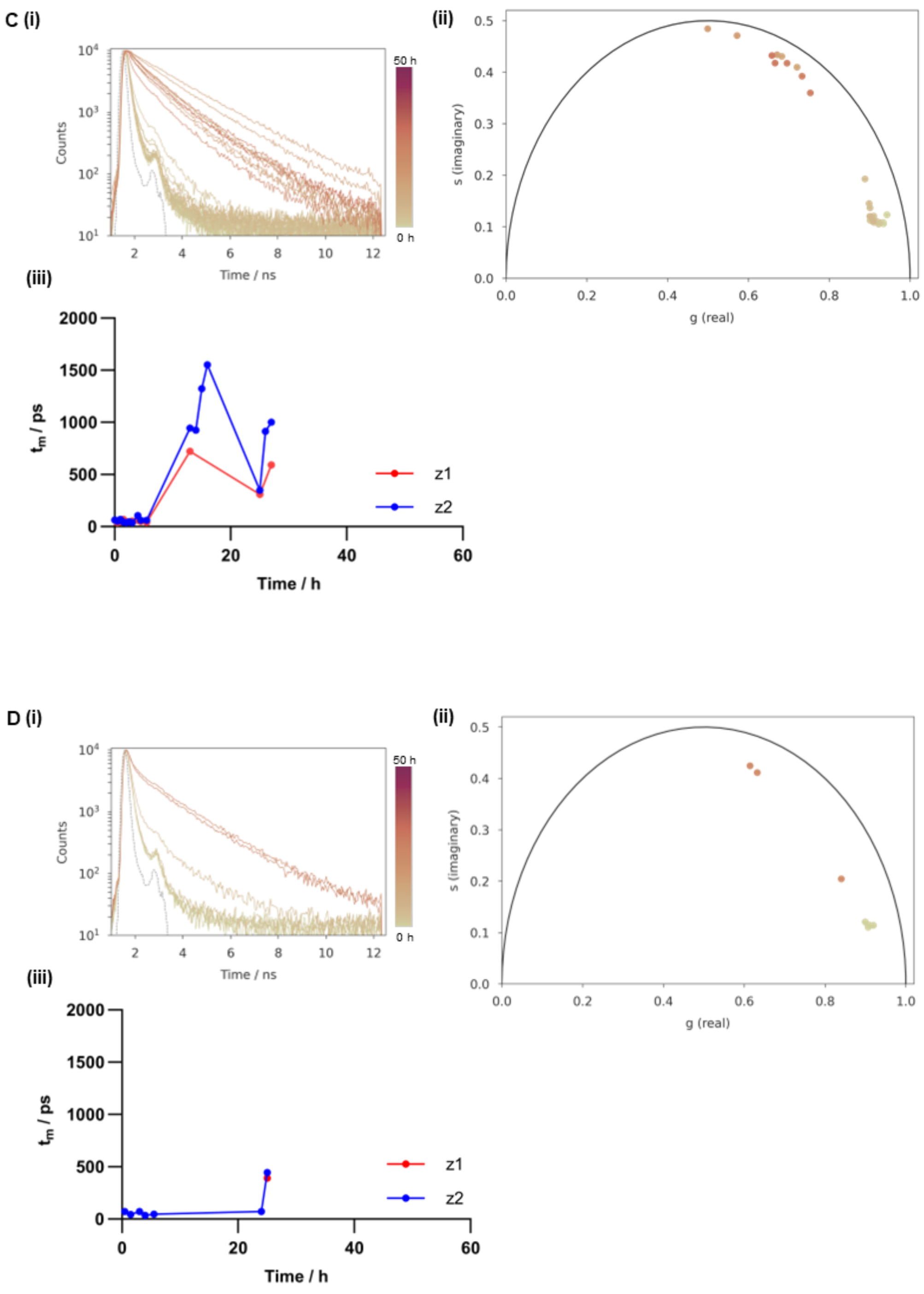


**Supplementary Fig. S4 Multiple repeats of $DiSC_2$ time-resolved fluorescence decays and lifetime analysis used to monitor WT $\alpha$Syn aggregation in PBS (pH 7.4) (A-D). (i)** Time-resolved fluorescence decays of $DiSC_2$ (3 $\mu$M) in the presence of aggregating WT $\alpha$Syn (150 $\mu$M). **(ii)** Phasor analysis of $DiSC_2$ decay profile. **(iii)** Fitted lifetimes ($\tau_m$) of $DiSC_2$ during the aggregation. z-positions were taken at 100 nm above the well plate surface (z2) and 1000 nm above this position (z1).

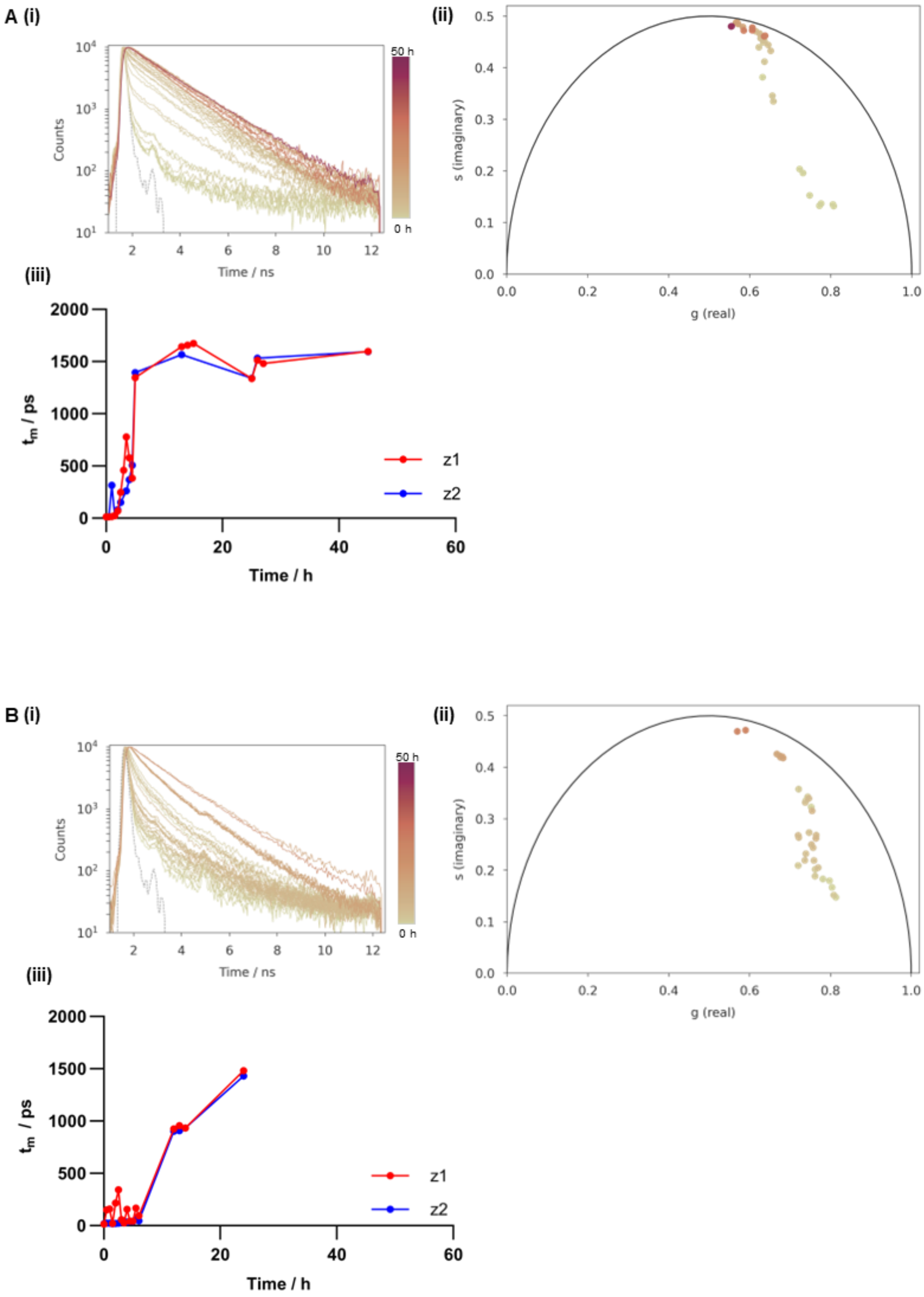

A (i)
Counts
Time / ns
50 h
0 h
(ii)
s (imaginary)
g (real)
(iii)
$t_m$ / ps
Time / h
z1
z2
B (i)
Counts
Time / ns
50 h
0 h
(ii)
s (imaginary)
g (real)
(iii)
$t_m$ / ps
Time / h
z1
z2

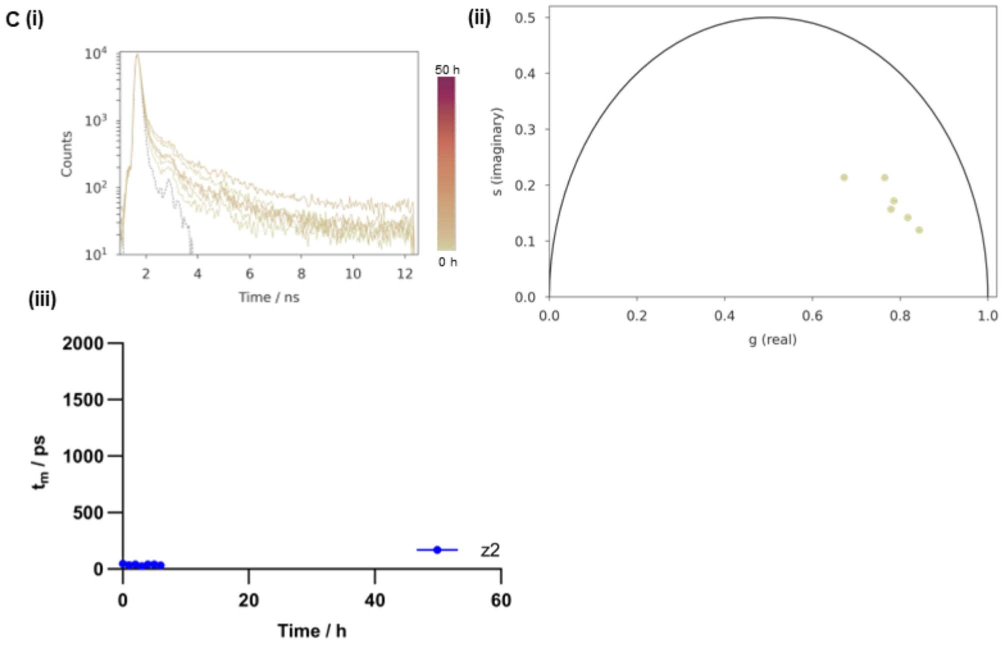
C (i)
(ii)
(iii)
Counts
Time / ns
50 h
0 h
s (imaginary)
g (real)
t_m / ps
Time / h
z2

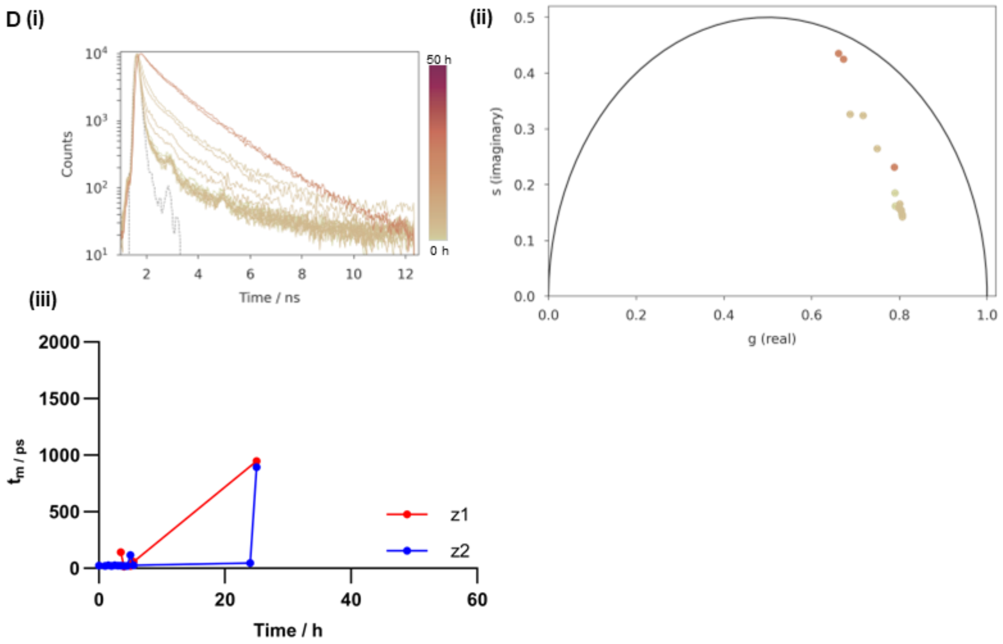
D (i)
(ii)
(iii)
Counts
Time / ns
50 h
0 h
s (imaginary)
g (real)
t_m / ps
Time / h
z1
z2

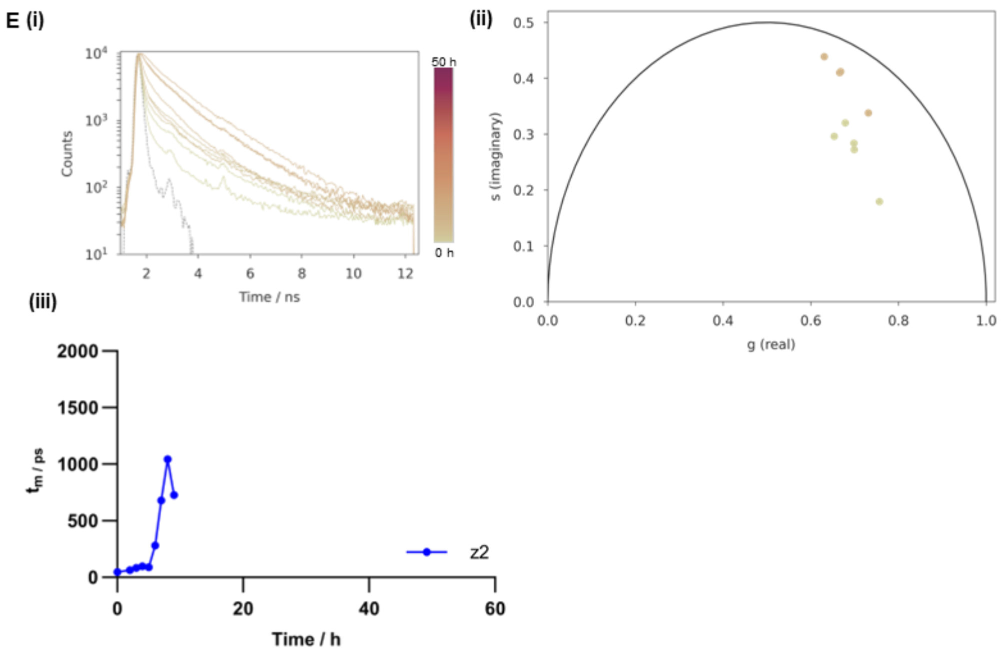


**Supplementary Fig. S5 Multiple repeats of ThT time-resolved fluorescence decays and lifetime analysis used to monitor WT $\alpha$Syn aggregation in PBS (pH 7.4) (A-E). (i)** Time-resolved fluorescence decays of ThT (10 $\mu$M) in the presence of aggregating WT $\alpha$Syn (150 $\mu$M). **(ii)** Phasor analysis of ThT decay profile. **(iii)** Fitted lifetimes ($\tau_m$) of ThT during the aggregation. z-positions were taken at 100 nm above the well plate surface (z2) and 1000 nm above this position (z1).

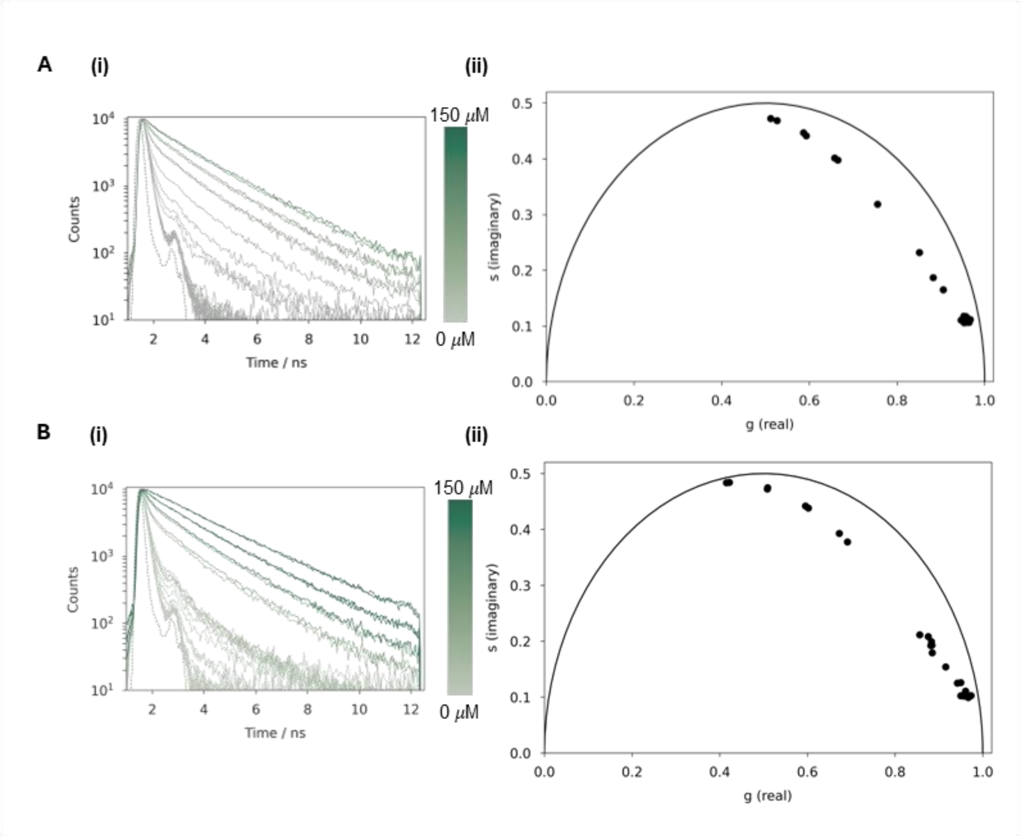


**Supplementary Fig. S6 Fluorescence lifetime analysis of $DiSC_2$ in the presence of isolated WT αSyn fibrils in PBS (pH 7.4) (A-B). (i)** Time-resolved fluorescence decays of $DiSC_2$ (3 μM) in the presence of increasing fibril concentration (0.001 μM – 150 μM) **(ii)** Phasor analysis of $DiSC_2$ decay profile.

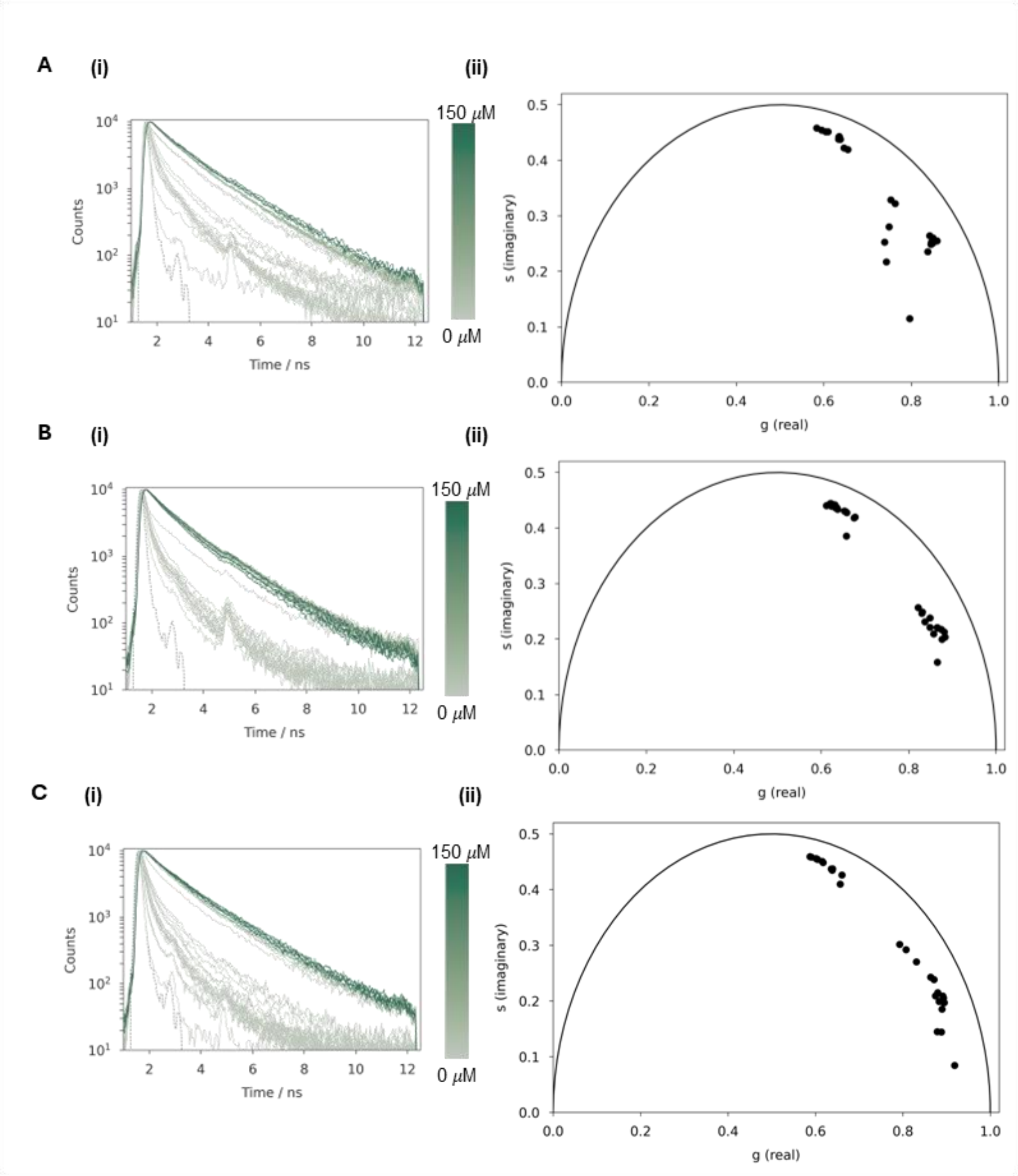


**Supplementary Fig. S7 Fluorescence lifetime analysis of ThT in the presence of isolated WT $\alpha$Syn fibrils in PBS (pH 7.4) (A-C). (i)** Time-resolved fluorescence decays of ThT (10 $\mu$M) in the presence of increasing fibril concentration (0.001 $\mu$M – 150 $\mu$M) **(ii)** Phasor analysis of ThT decay profile.

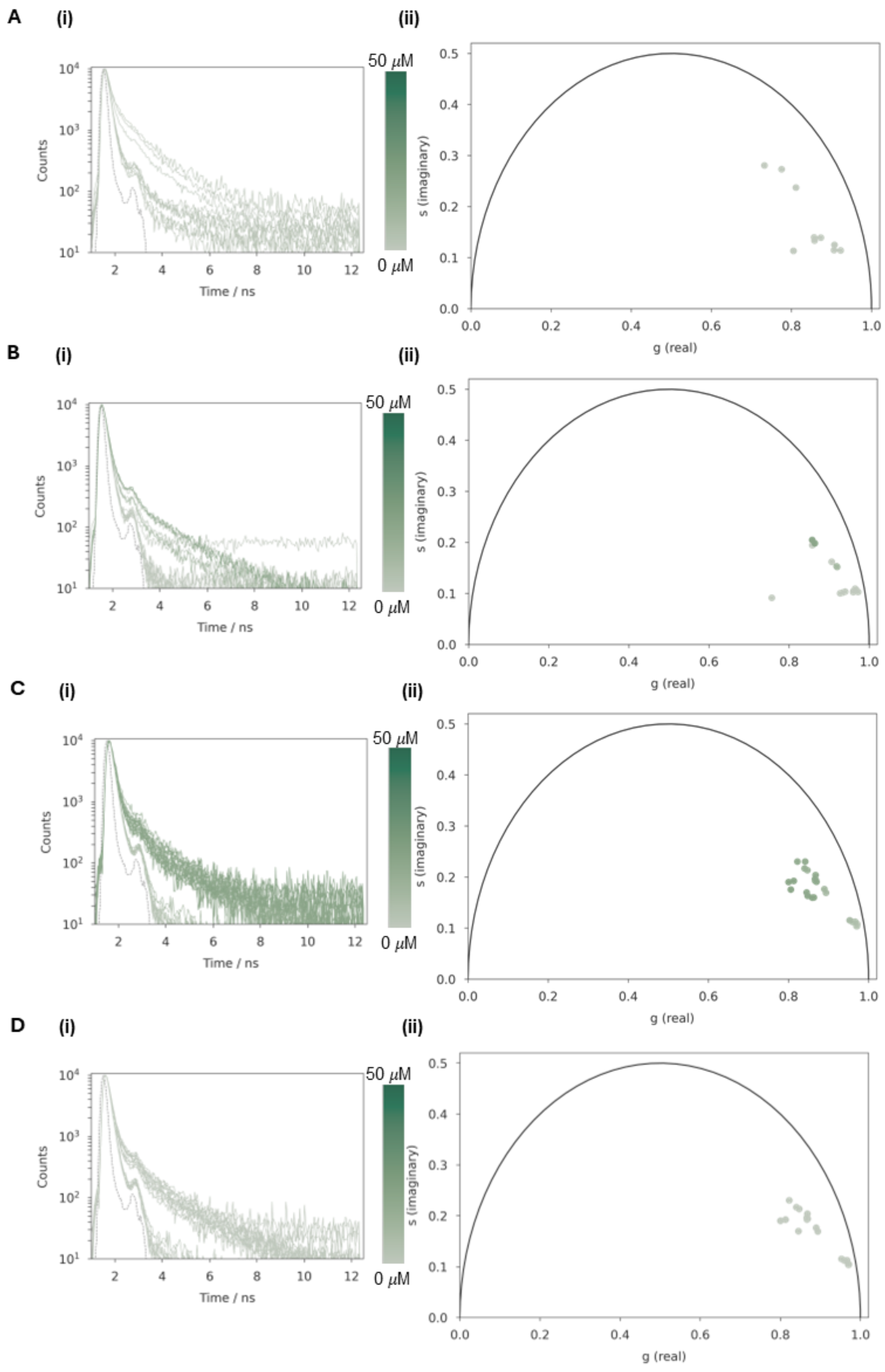


**Supplementary Fig. S8 Fluorescence lifetime analysis of $DiSC_2$ in the presence of isolated WT $\alpha$Syn stabilized oligomers in PBS (pH 7.4) (A-D). (i)** Time-resolved fluorescence decays of $DiSC_2$ (3 $\mu$M) in the presence of increasing stabilized oligomer concentration (0.001 $\mu$M – 20 $\mu$M) **(ii)** Phasor analysis of $DiSC_2$ decay profile.

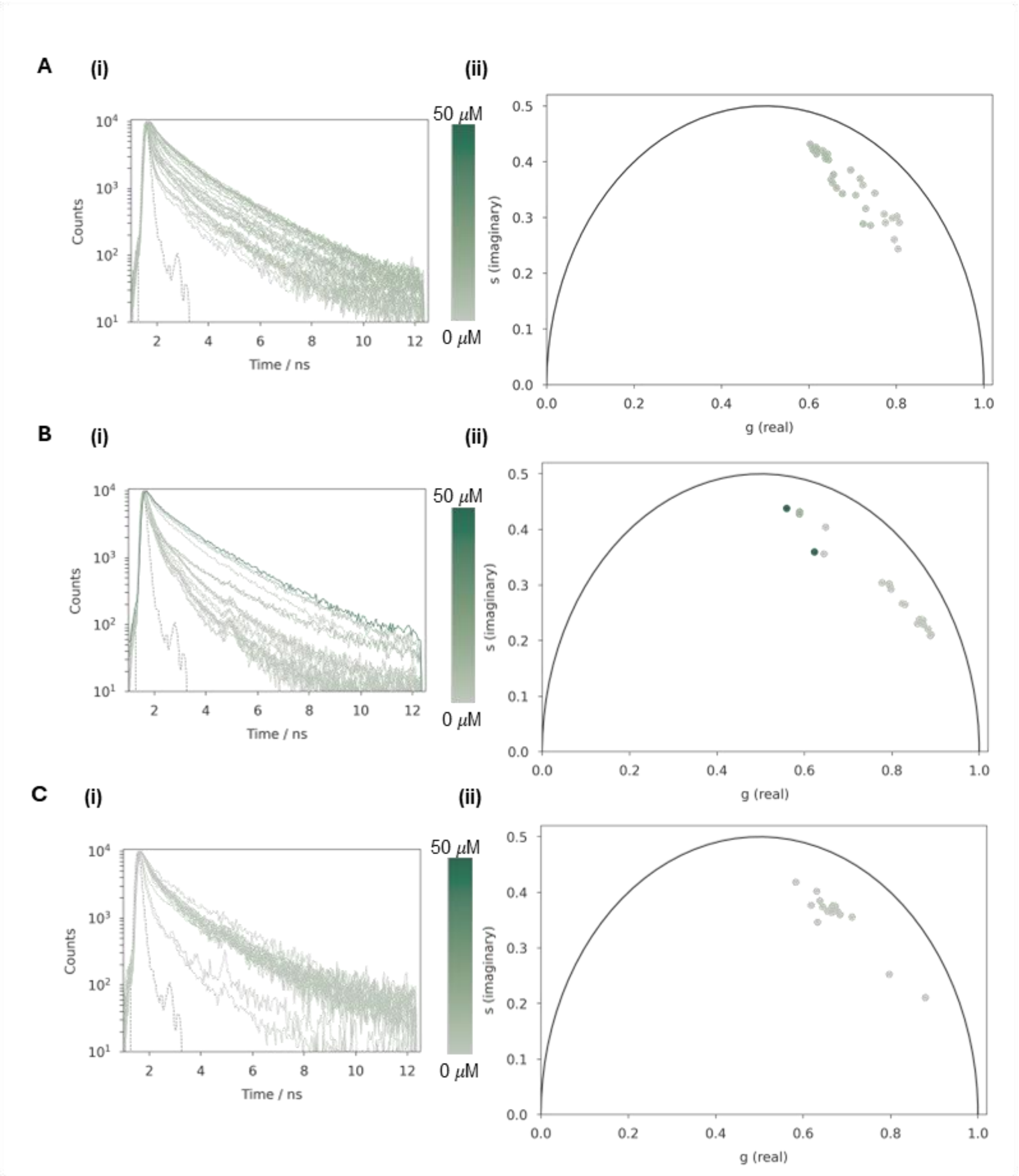


**Supplementary Fig. S9 Fluorescence lifetime analysis of ThT in the presence of isolated WT $\alpha$Syn stabilized oligomers in PBS (pH 7.4) (A-C). (i)** Time-resolved fluorescence decays of ThT (10 $\mu$M) in the presence of increasing stabilized oligomer concentration (0.001 $\mu$M – 50 $\mu$M) **(ii)** Phasor analysis of ThT decay profile.

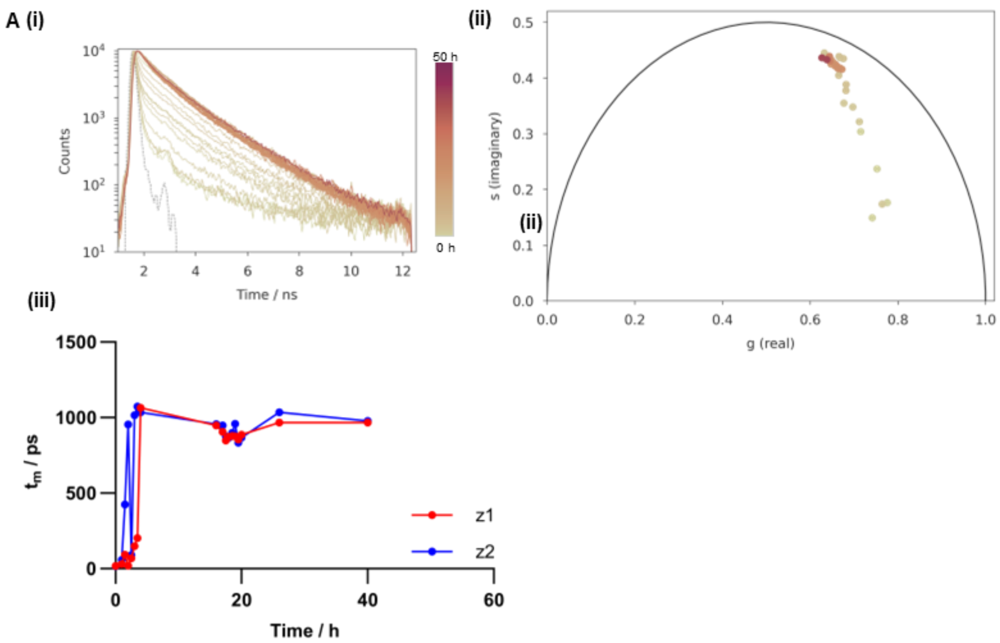
A (i)
(ii)
(iii)
Counts
Time / ns
50 h
0 h
s (imaginary)
g (real)
tm / ps
Time / h
z1
z2

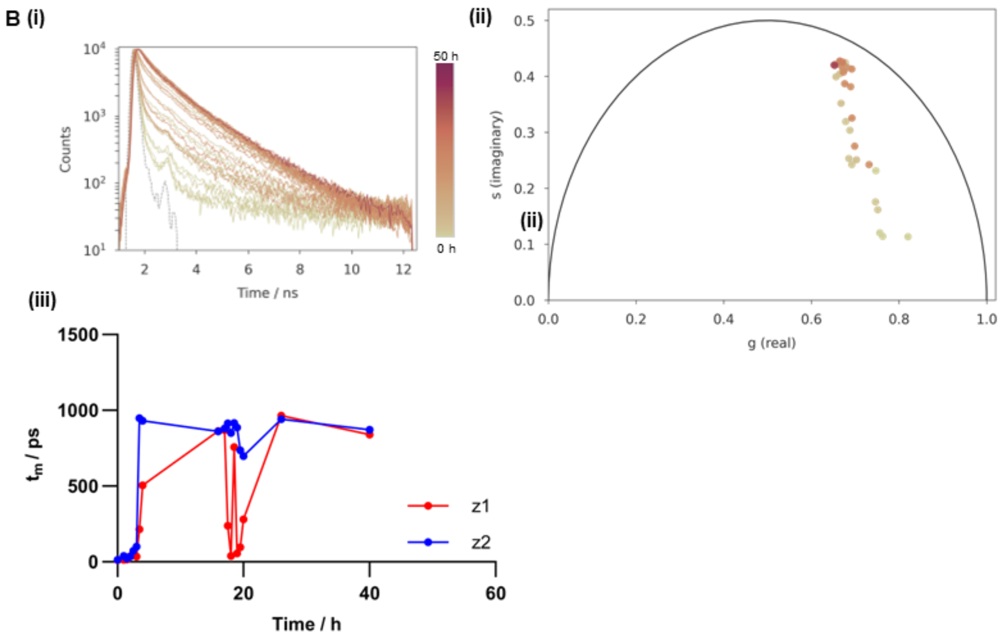
B (i)
(ii)
(iii)
Counts
Time / ns
50 h
0 h
s (imaginary)
g (real)
tm / ps
Time / h
z1
z2

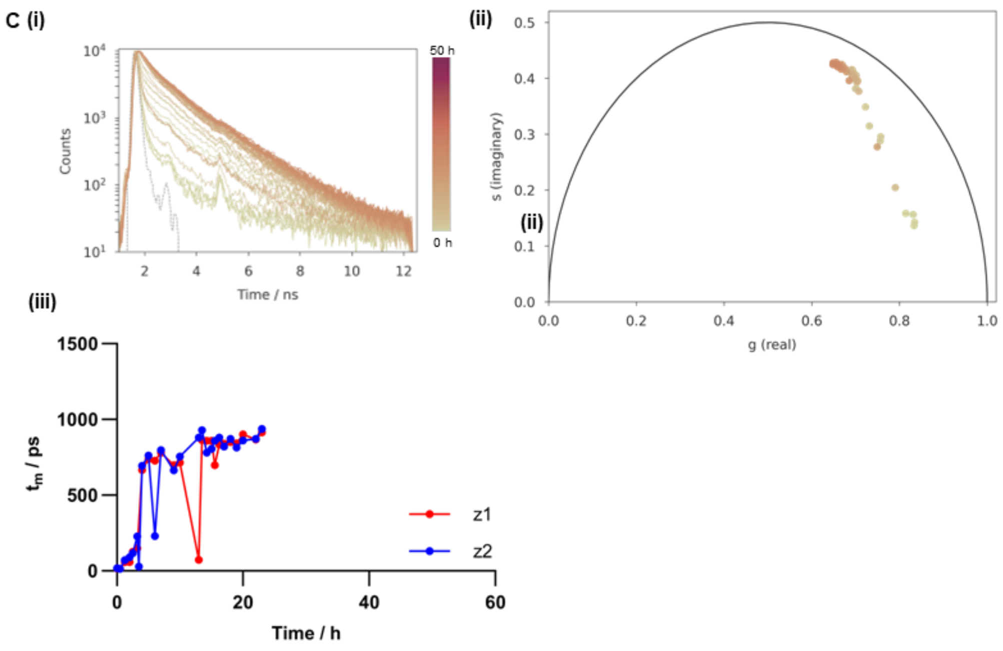


**Supplementary Fig. S10 Multiple repeats of ThT time-resolved fluorescence decays and lifetime analysis used to monitor A30P $\alpha$Syn aggregation in PBS (pH 7.4) (A-C). (i)** Time-resolved fluorescence decays of ThT (10 $\mu$M) in the presence of aggregating A30P $\alpha$Syn (150 $\mu$M). **(ii)** Phasor analysis of ThT decay profile. **(iii)** Fitted lifetimes ($\tau_m$) of ThT during the aggregation. z-positions were taken at 100 nm above the well plate surface (z2) and 1000 nm above this position (z1).

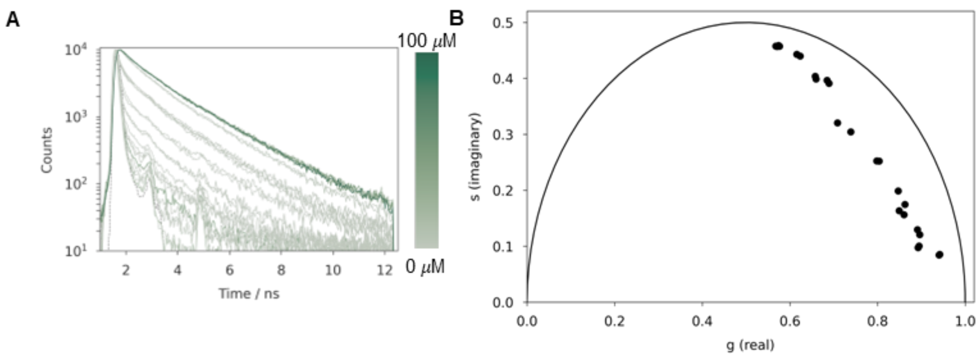


**Supplementary Fig. S11 Fluorescence lifetime analysis of ThT in the presence of isolated A30P $\alpha$Syn fibrils in PBS (pH 7.4) (A-C). (i)** Time-resolved fluorescence decays of ThT (10 $\mu$M) in the presence of increasing fibril concentration (0.001 $\mu$M – 150 $\mu$M) **(ii)** Phasor analysis of ThT decay profile.

A

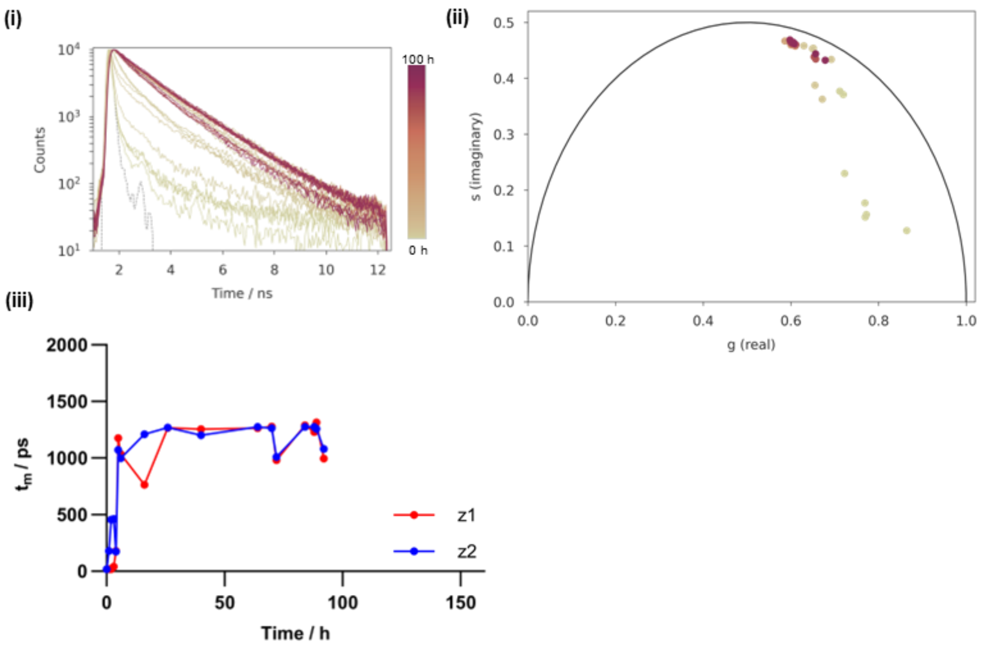
(i)
10⁴
10³
10²
10¹
Counts
2 4 6 8 10 12
Time / ns
100 h
0 h
(ii)
0.5
0.4
0.3
0.2
0.1
0.0
s (imaginary)
0.0 0.2 0.4 0.6 0.8 1.0
g (real)
(iii)
2000
1500
1000
500
0
t_m / ps
0 50 100 150
Time / h
z1
z2

B

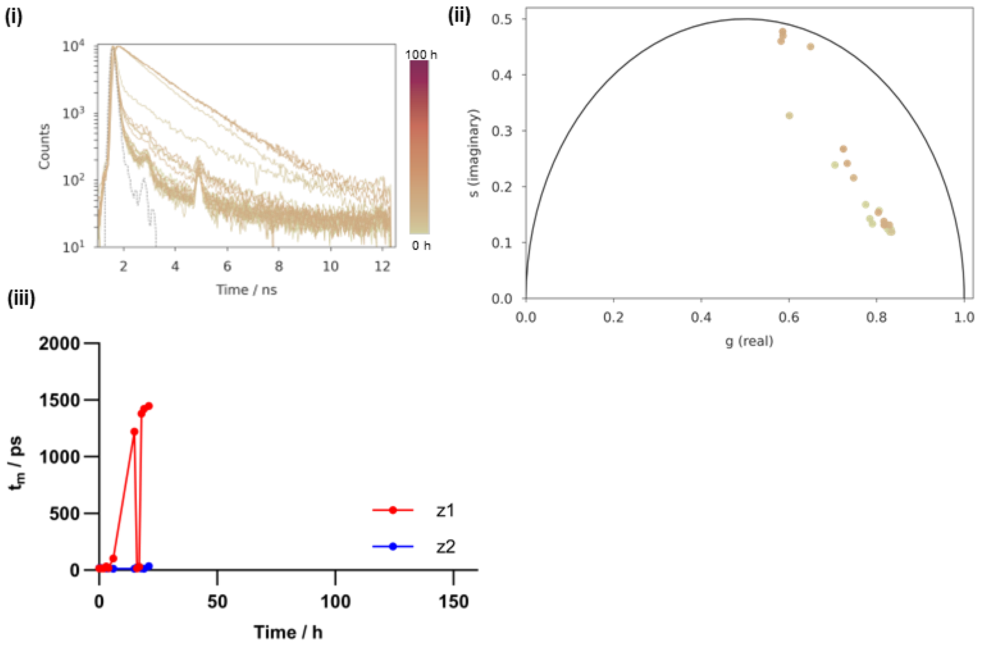
(i)
10⁴
10³
10²
10¹
Counts
2 4 6 8 10 12
Time / ns
100 h
0 h
(ii)
0.5
0.4
0.3
0.2
0.1
0.0
s (imaginary)
0.0 0.2 0.4 0.6 0.8 1.0
g (real)
(iii)
2000
1500
1000
500
0
t_m / ps
0 50 100 150
Time / h
z1
z2

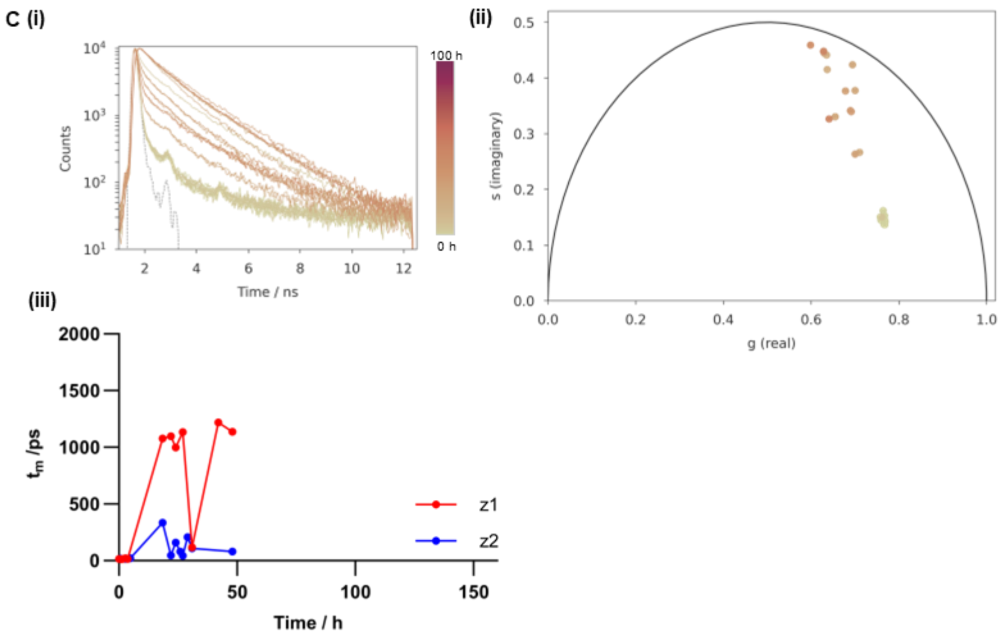


**Supplementary Fig. S12 Multiple repeats of ThT time-resolved fluorescence decays and lifetime analysis used to monitor WT $\alpha$Syn aggregation in 20 mM Tris-HCl with 200 mM NaCl (pH 7.5) (A-B). (i)** Time-resolved fluorescence decays of ThT (10 $\mu$M) in the presence of aggregating WT $\alpha$Syn (100 $\mu$M). **(ii)** Phasor analysis of ThT decay profile. **(iii)** Fitted lifetimes ($\tau_m$) of ThT during the aggregation. z-positions were taken at 100 nm above the well plate surface (z2) and 1000 nm above this position (or until the sample surface was reached) (z1).

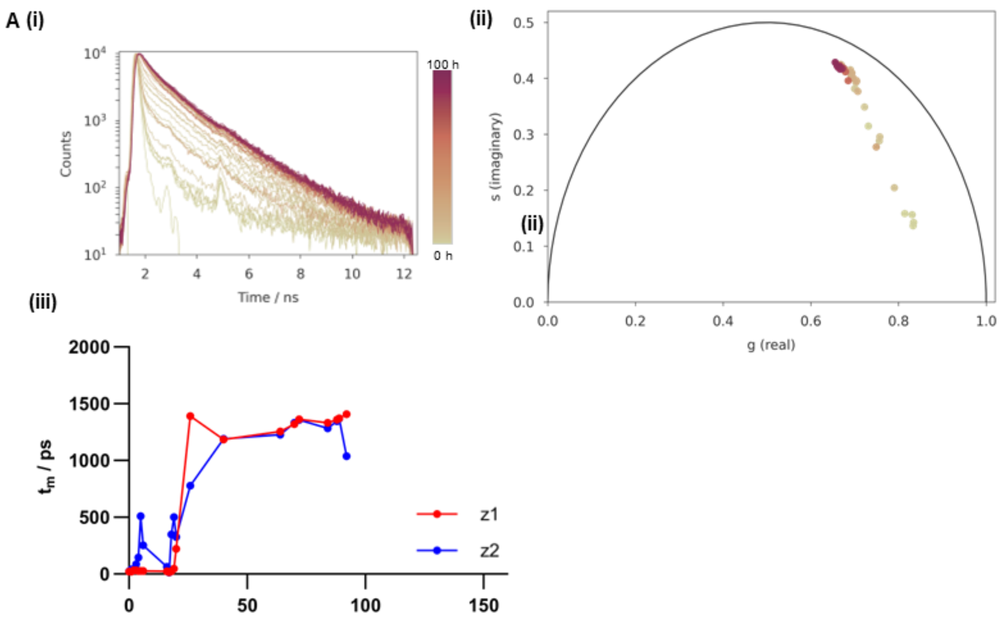
A (i)
(ii)
(iii)
Counts
Time / ns
100 h
0 h
s (imaginary)
g (real)
(ii)
$t_m$ / ps
z1
z2

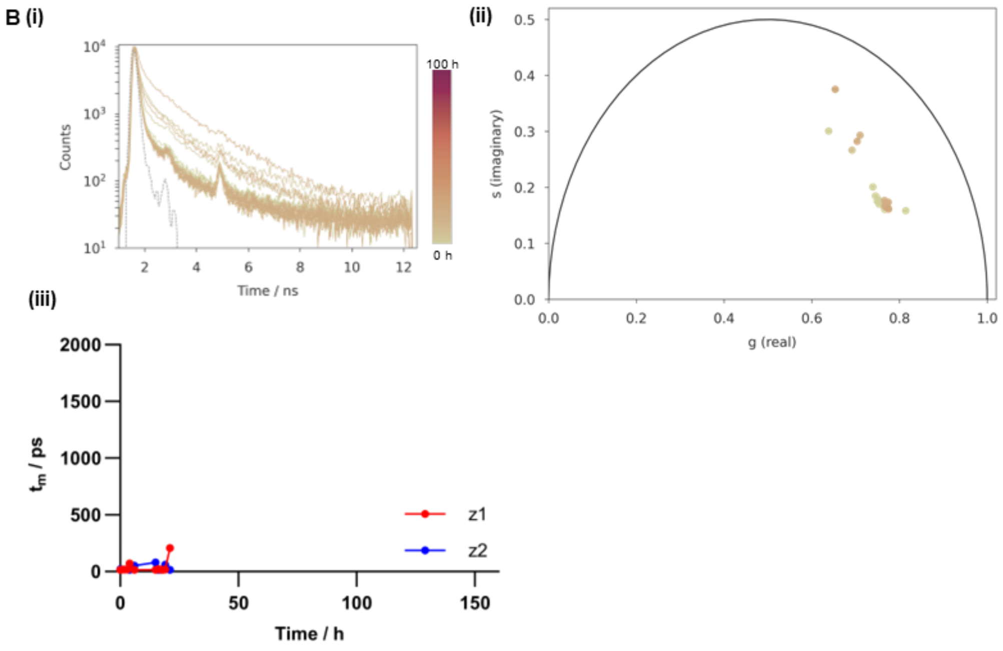
B (i)
(ii)
(iii)
Counts
Time / ns
100 h
0 h
s (imaginary)
g (real)
$t_m$ / ps
z1
z2
Time / h

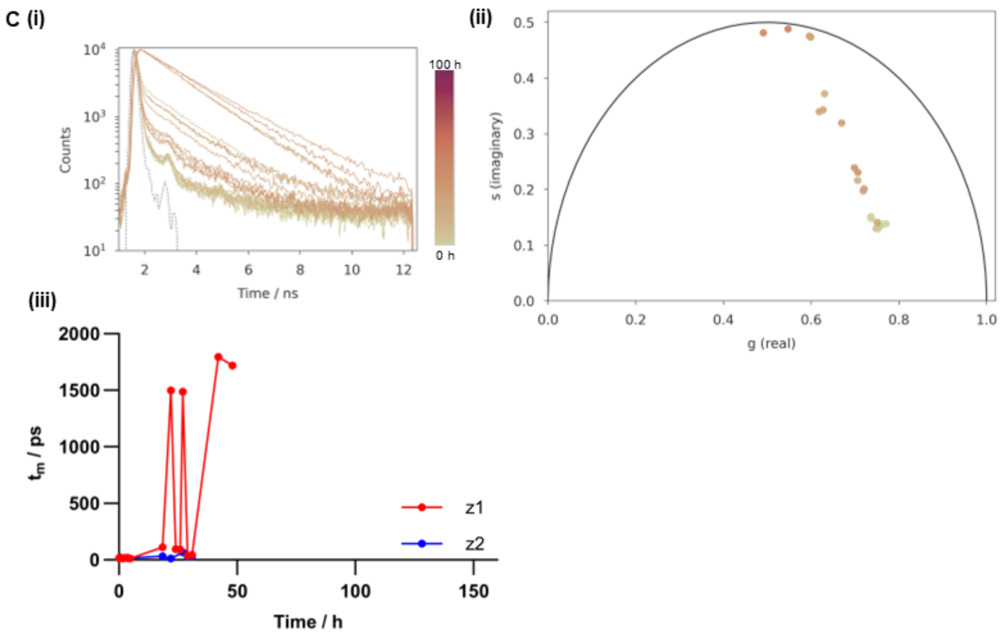


**Supplementary Fig. S13 Multiple repeats of ThT time-resolved fluorescence decays and lifetime analysis used to monitor ΔP1 $\alpha$Syn aggregation in 20 mM Tris-HCl with 200 mM NaCl (pH 7.5) (A-B). (i)** Time-resolved fluorescence decays of ThT (10 $\mu$M) in the presence of aggregating ΔP1 $\alpha$Syn (100 $\mu$M). **(ii)** Phasor analysis of ThT decay profile. **(iii)** Fitted lifetimes ($\tau_m$) of ThT during the aggregation. z-positions were taken at 100 nm above the well plate surface (z2) and 1000 nm above this position (or until the sample surface was reached) (z1).

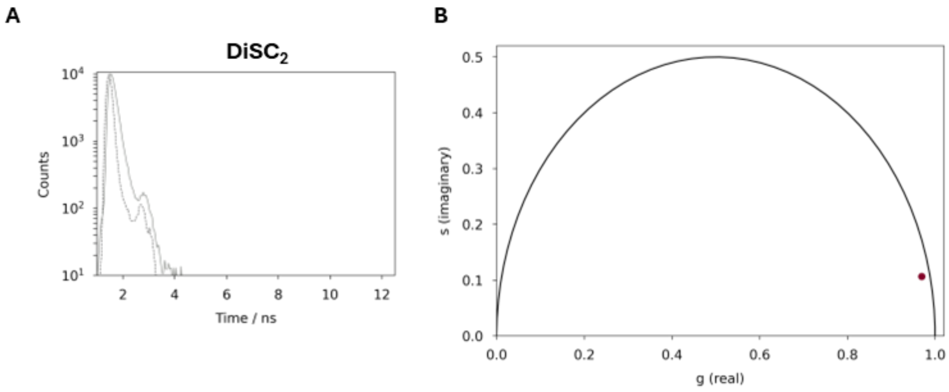


**Supplementary Fig. S14 Fluorescence lifetime analysis of free DiSC$_2$ in PBS (pH 7.4) (A-B). A** Time-resolved fluorescence decay of free DiSC$_2$ (3 $\mu$M). **B** Phasor analysis of DiSC$_2$ decay profile.

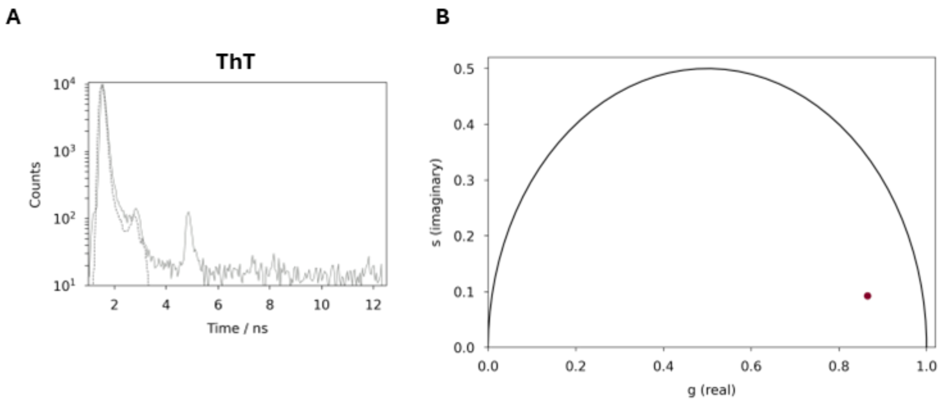


**Supplementary Fig. S15 Fluorescence lifetime analysis of free ThT in PBS (pH 7.4) (A-B). A** Time-resolved fluorescence decay of free ThT (10 $\mu$M). **B** Phasor analysis of ThT decay profile.

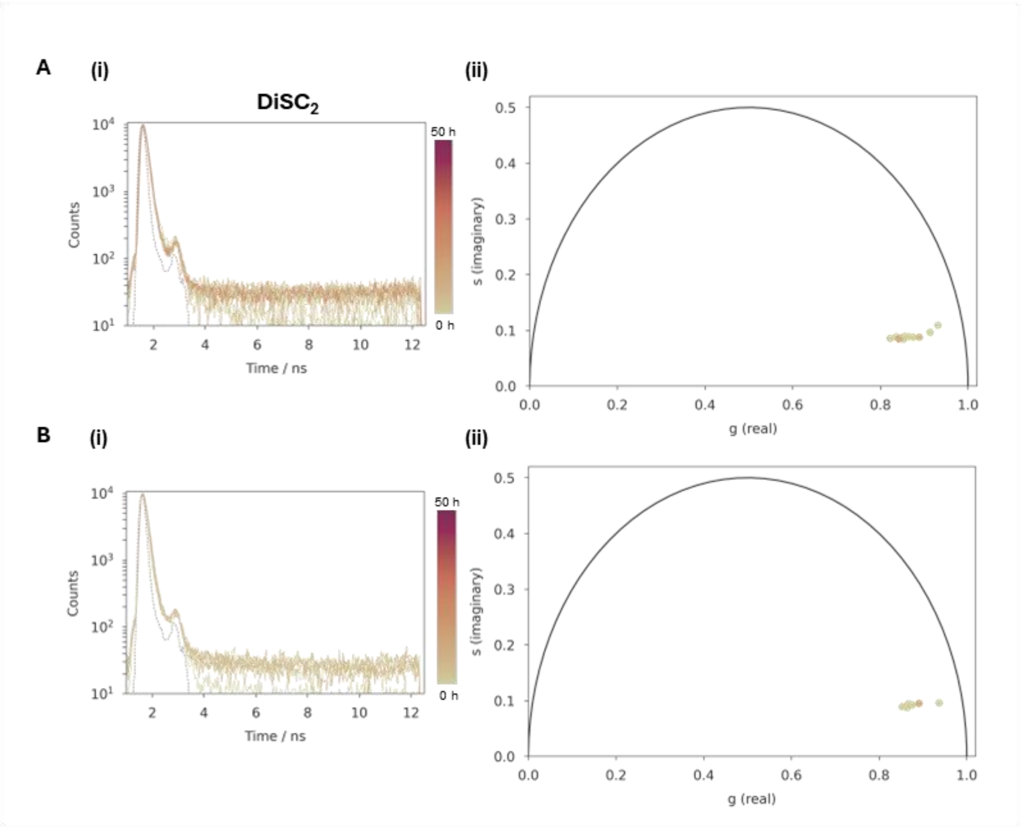


**Supplementary Fig. S16 DiSC$_2$ time-resolved fluorescence decays and phasor analysis in PBS (pH 7.4) (A-B). (i)** Time-resolved fluorescence decays of DiSC$_2$ (3 $\mu$M) in PBS (pH 7.4) over 27 h (A) and 24 h (B) . **(ii)** Phasor analysis of DiSC$_2$ decay profile.

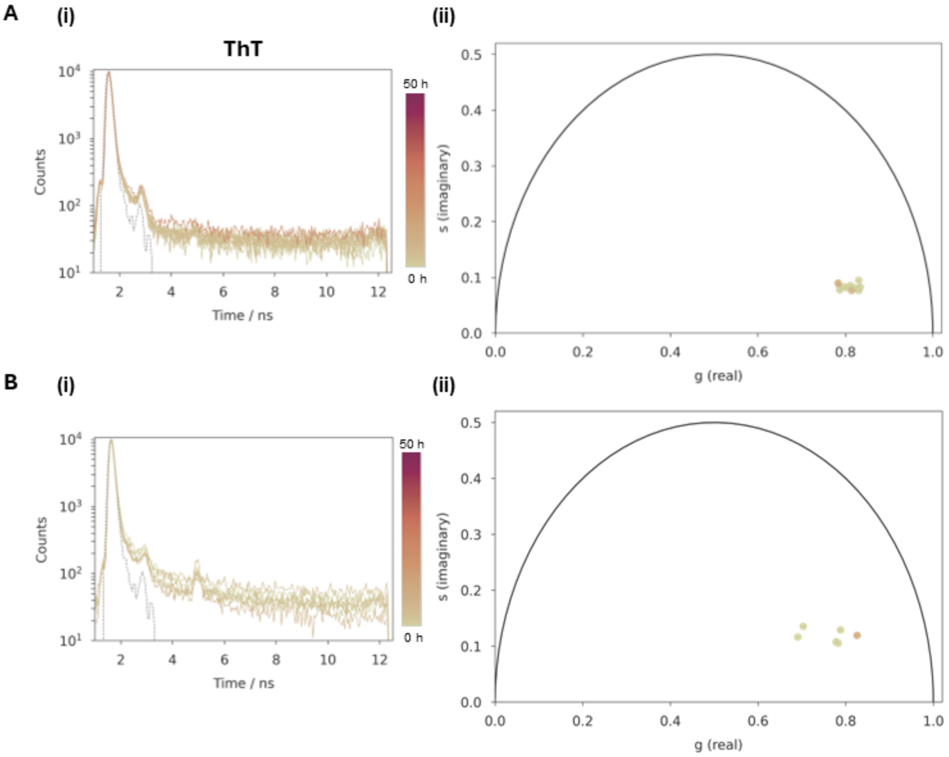


**Supplementary Fig. S17 ThT time-resolved fluorescence decays and phasor analysis in PBS (pH 7.4) (A-B). (i)** Time-resolved fluorescence decays of ThT (10 $\mu$M) in PBS (pH 7.4) over 27 h (A) and 24 h (B). **(ii)** Phasor analysis of ThT decay profile.

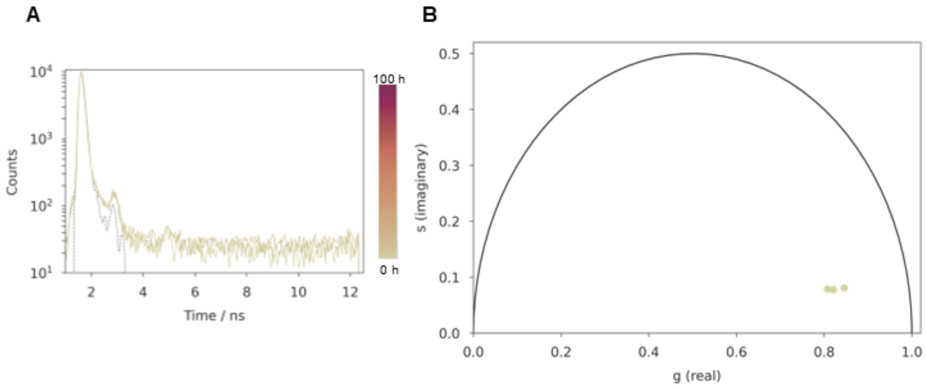


**Supplementary Fig. S18 ThT time-resolved fluorescence decays and phasor analysis in 20 mM Tris-HCl with 200 mM NaCl (pH 7.5).** **A** Time-resolved fluorescence decays of ThT (10 $\mu$M) in 20 mM Tris-HCl with 200 mM NaCl (pH 7.5). **B** Phasor analysis of ThT decay profile.

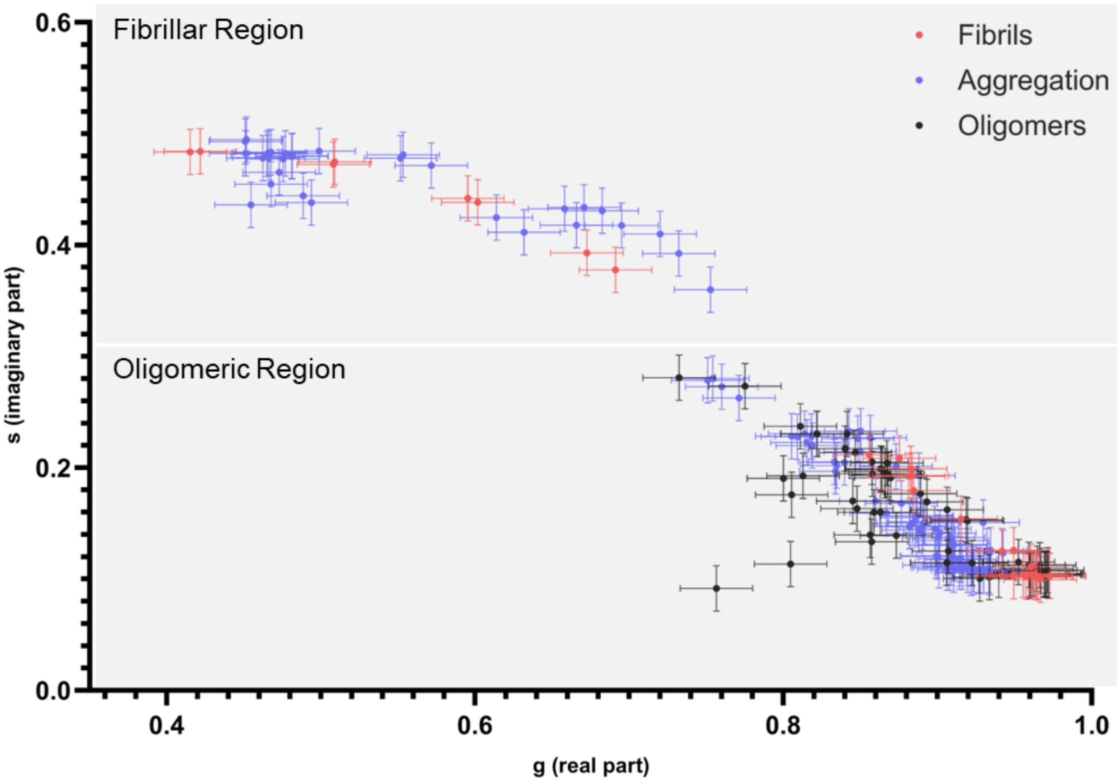


**Supplementary Fig. S19 Classification of the separate regions of the DiSC$_2$ phasor plot.** Error bars represent the standard deviation ($2\sigma$) of technical repeats of FLIM fibril and oligomer measurements. Two points are considered overlapping if there is overlap in two error bar directions. Regions are assigned if there is at least a 60 % overlap between points. In this case ~70 % of aggregation points in the fibrillar region overlap with the fibrillar points. In the oligomeric region, ~60 % of the aggregation points deviate from the fibrillar region and all the stabilized oligomers reside in this region, with all the aggregation data points overlapping with the stabilized oligomers.

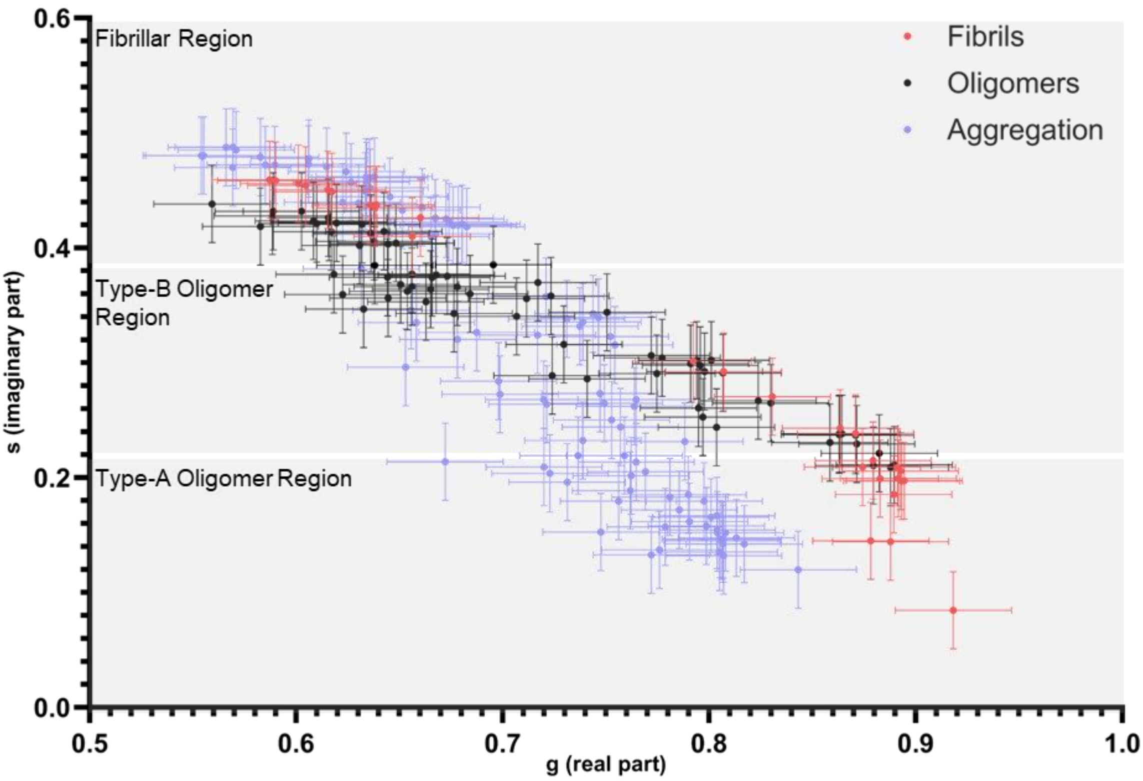


**Supplementary Fig. S20 Classification of the separate regions of the ThT phasor plot.** Error bars represent the standard deviation ($2\sigma$) of technical repeats of FLIM fibril and oligomer measurements. Two points are considered overlapping if there is overlap in two error bar directions. Regions are assigned if there is at least a 60 % overlap between points. In this case ~94 % of aggregation points in the fibrillar region overlap with the fibrillar points. In the Type-B oligomer region ~69 % aggregation points overlap with the stabilized oligomer points and in the Type-A region there is less than 4 % overlap of the aggregation data with either fibrillar or stabilized oligomer points.

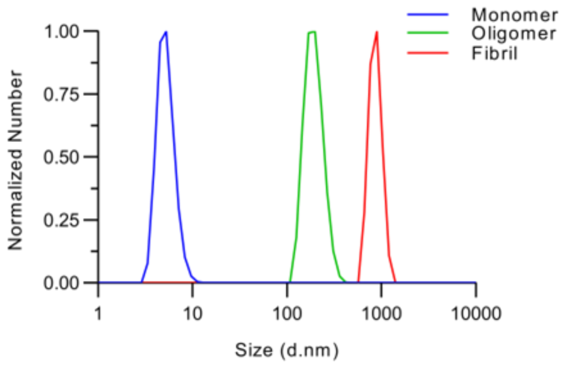


**Supplementary Fig. S21 Dynamic light scattering (DLS) analysis of monomeric (blue), oligomeric (green) and fibrillar WT $\alpha$Syn (red).**

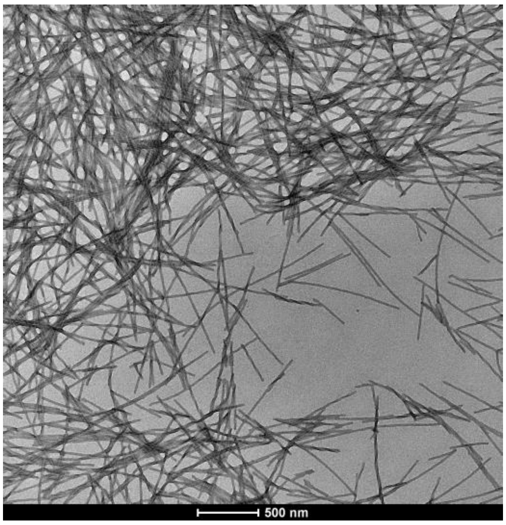


**Supplementary Fig. S22 Representative electron microscopy (EM) image of fully formed WT $\alpha$Syn fibrils (150 $\mu$M).**

**WT Sian_31030** **29-May-2024**

WT Sian_31030 241 (1.514) Cm (227:276) 1: TOF MS ES+ 805

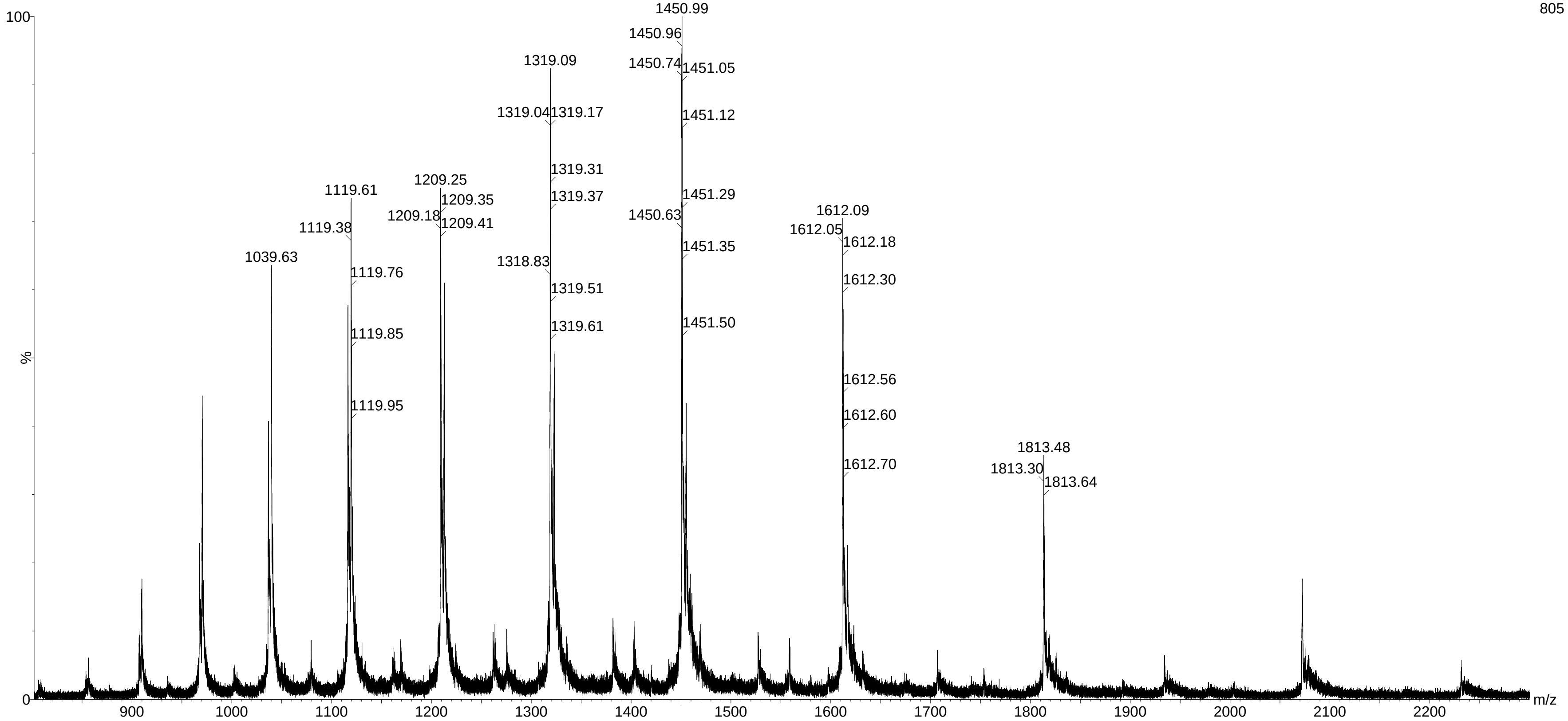

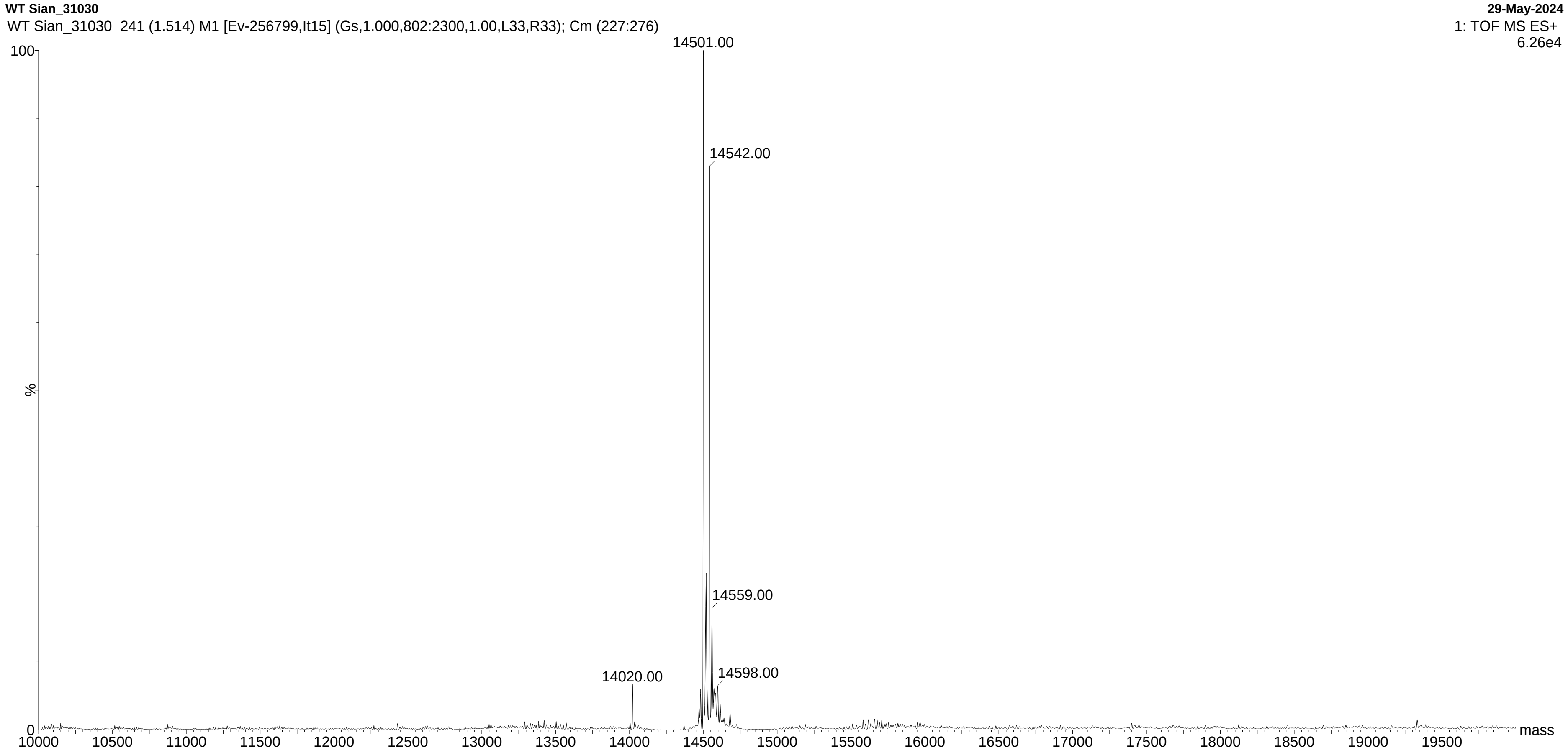


**Supplementary Fig. S23 Time-of-flight (TOF) mass spectrometry (MS) electrospray ionization (ES+) Mass spectrometry spectrum of pure WT $\alpha$Syn in water.**

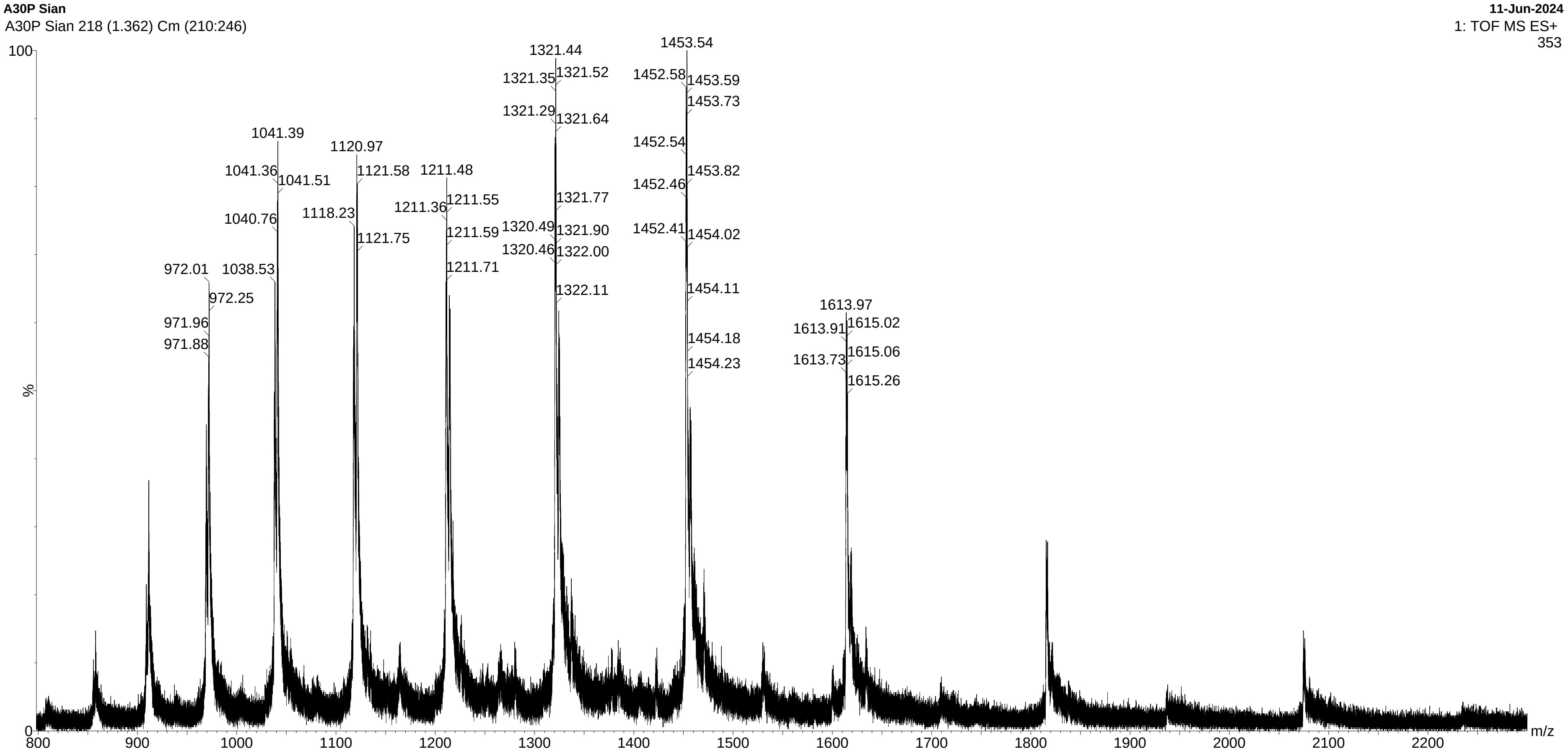
A30P Sian
A30P Sian 218 (1.362) Cm (210:246)
11-Jun-2024
1: TOF MS ES+
353
100
%
0
m/z
800
900
1000
1100
1200
1300
1400
1500
1600
1700
1800
1900
2000
2100
2200
972.01
972.25
971.96
971.88
1041.39
1041.36
1041.51
1040.76
1038.53
1120.97
1121.58
1118.23
1121.75
1211.48
1211.55
1211.36
1211.59
1211.71
1321.44
1321.35
1321.52
1321.29
1321.64
1321.77
1320.49
1321.90
1320.46
1322.00
1322.11
1453.54
1452.58
1453.59
1453.73
1452.54
1453.82
1452.46
1452.41
1454.02
1454.11
1454.18
1454.23
1613.97
1613.91
1615.02
1613.73
1615.06
1615.26

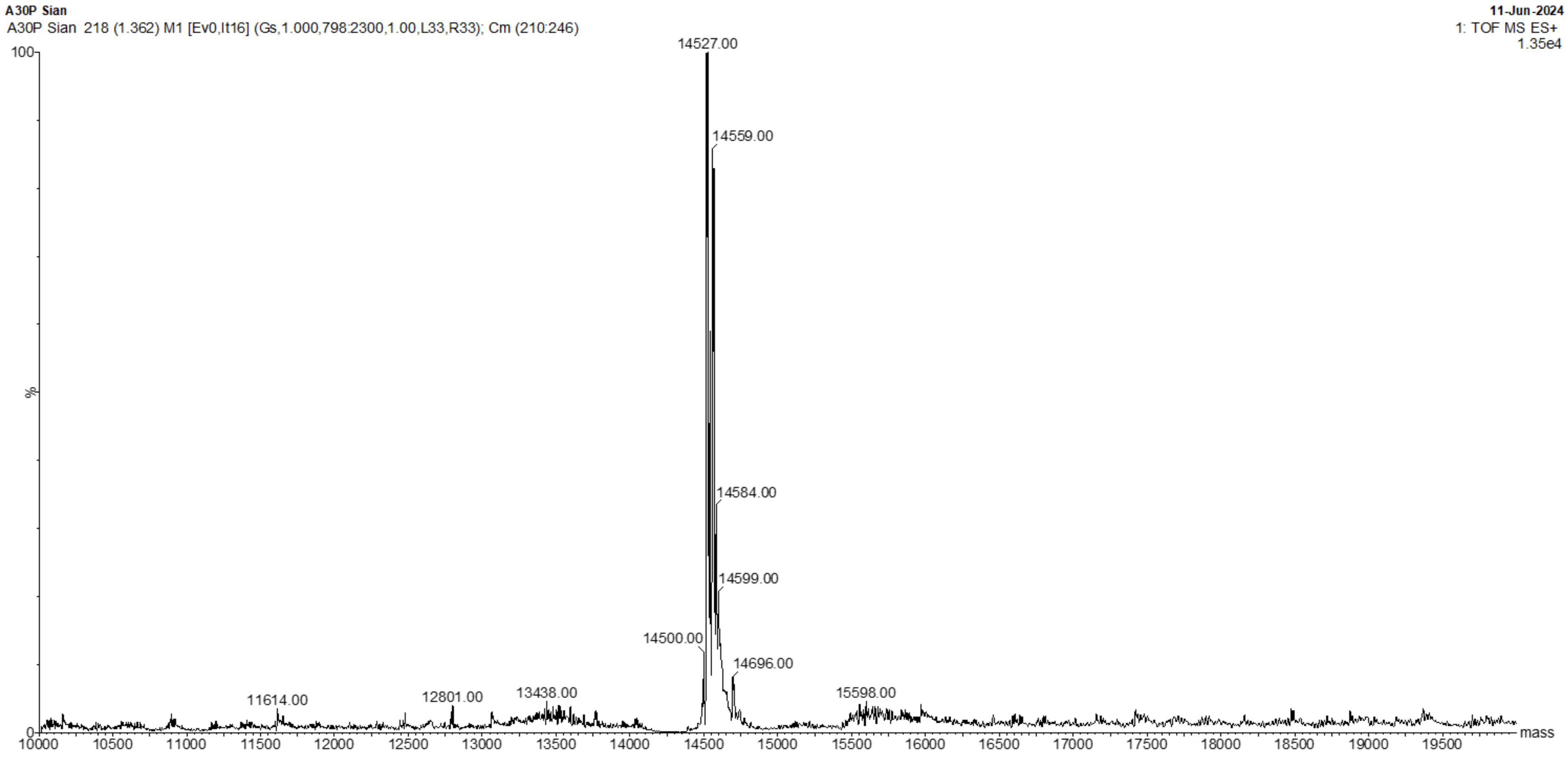


**Supplementary Fig. S24 TOF MS ES+ of pure A30P $\alpha$Syn in water.**

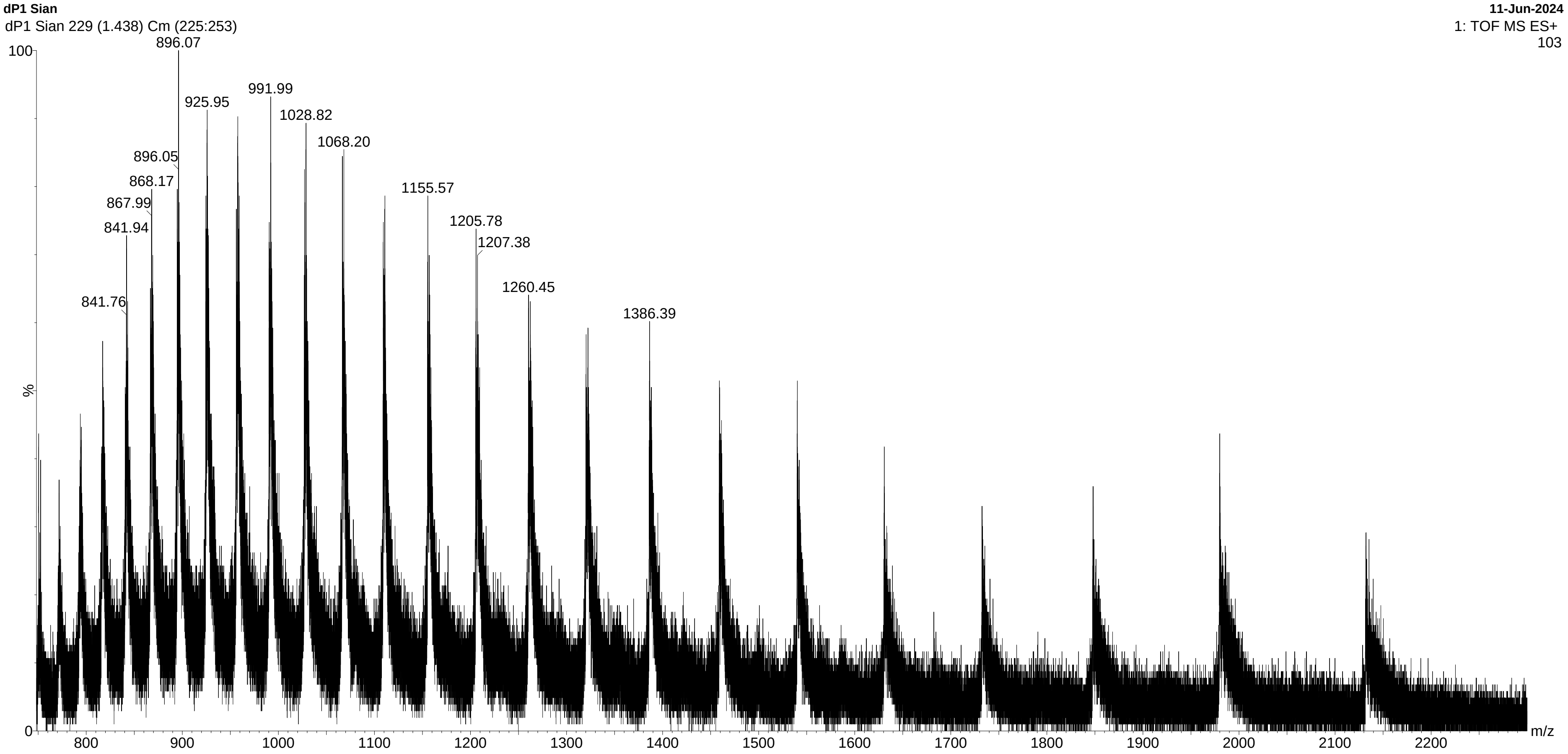
dP1 Sian
dP1 Sian 229 (1.438) Cm (225:253)
11-Jun-2024
1: TOF MS ES+
103
896.07
925.95
991.99
1028.82
1068.20
896.05
868.17
867.99
841.94
1155.57
1205.78
1207.38
1260.45
841.76
1386.39
100
%
0
800
900
1000
1100
1200
1300
1400
1500
1600
1700
1800
1900
2000
2100
2200
m/z

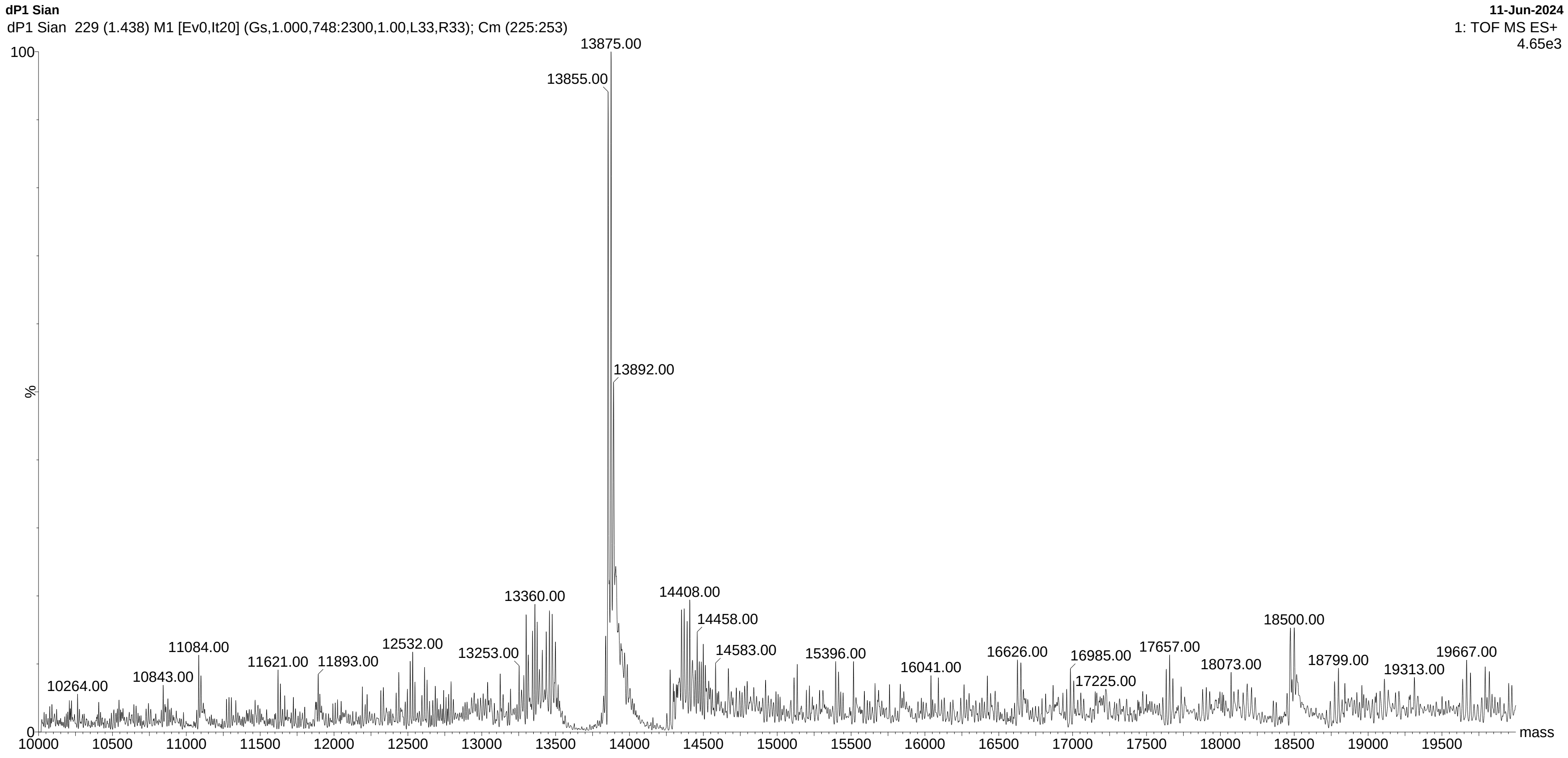


**Supplementary Fig. S25 TOF MS ES+ spectrum of pure ΔP1 αSyn in water.**

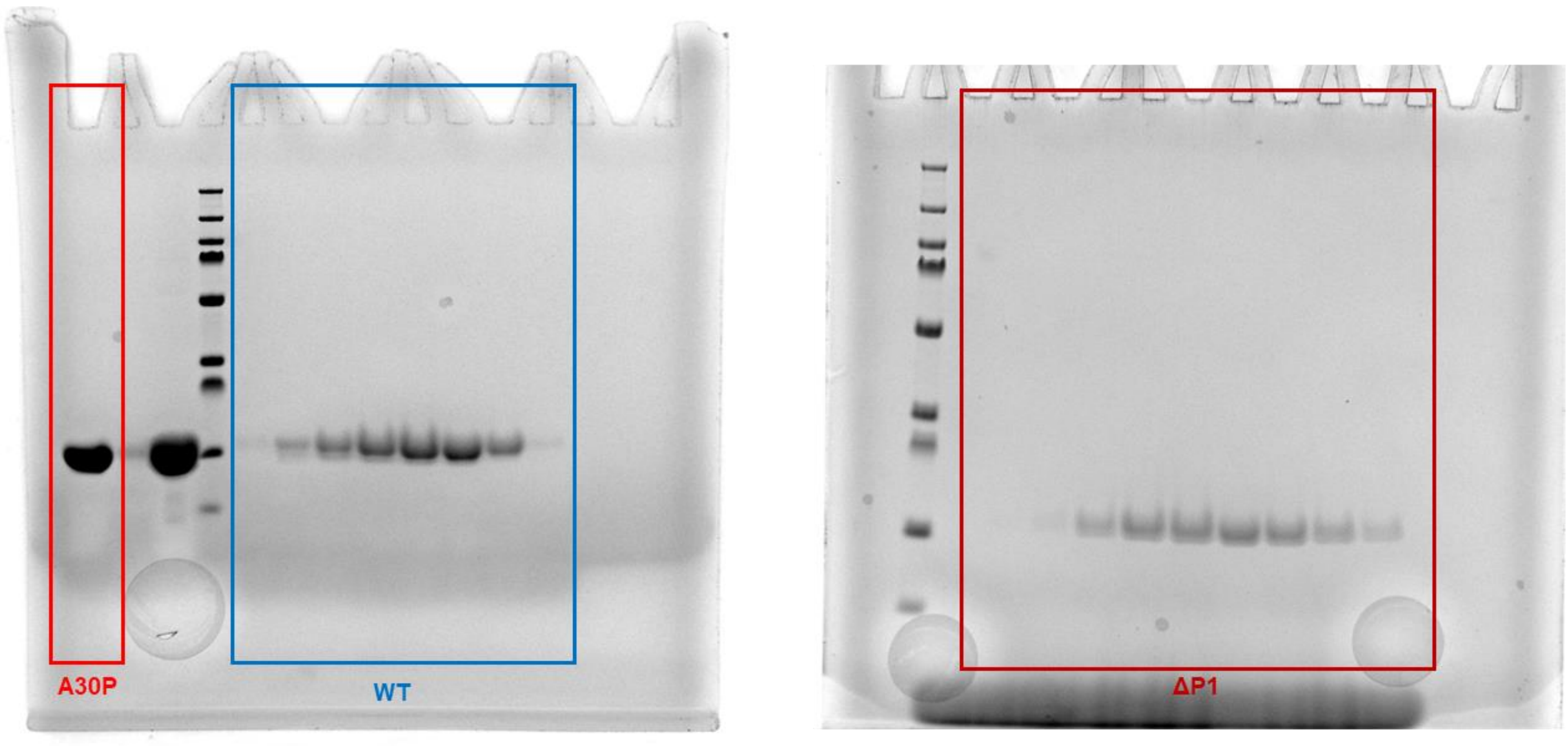


**Supplementary Fig. S26 Sodium dodecyl sulfate–polyacrylamide gel electrophoresis (SDS-PAGE) image of purified A30P, WT and ΔP1 $\alpha$Syn after SEC.** A30P $\alpha$Syn fractions from SEC have been combined. WT and ΔP1 $\alpha$Syn fractions from SEC have been imaged before being combined.
